# STARRPeaker: Uniform processing and accurate identification of STARR-seq active regions

**DOI:** 10.1101/694869

**Authors:** Donghoon Lee, Manman Shi, Jennifer Moran, Martha Wall, Jing Zhang, Jason Liu, Dominic Fitzgerald, Yasuhiro Kyono, Lijia Ma, Kevin P White, Mark Gerstein

## Abstract

**Background:** High-throughput reporter assays, such as self-transcribing active regulatory region sequencing (STARR-seq), allow for unbiased and quantitative assessment of enhancers at a genome-wide scale. Recent advances in STARR-seq technology have employed progressively more complex genomic libraries and increased sequencing depths, to assay larger sized regions, up to the entire human genome. These advances necessitate a reliable processing pipeline and peak-calling algorithm.

**Results:** Most STARR-seq studies have relied on chromatin immunoprecipitation sequencing (ChIP-seq) processing pipelines. However, there are key differences in STARR-seq versus ChIP-seq. First, STARR-seq uses transcribed RNA to measure the activity of an enhancer, making an accurate determination of the basal transcription rate important. Second, STARR-seq coverage is highly non-uniform, overdispersed, and often confounded by sequencing biases, such as GC content and mappability. Lastly, here, we observed a clear correlation between RNA thermodynamic stability and STARR-seq readout, suggesting that STARR-seq may be sensitive to RNA secondary structure and stability. Considering these findings, we developed a negative-binomial regression framework for uniformly processing STARR-seq data, called STARRPeaker. In support of this, we generated whole-genome STARR-seq data from the HepG2 and K562 human cell lines and applied STARRPeaker to call enhancers.

**Conclusions:** We show STARRPeaker can unbiasedly detect active enhancers from both captured and whole-genome STARR-seq data. Specifically, we report ∼33,000 and ∼20,000 candidate enhancers from HepG2 and K562, respectively. Moreover, we show that STARRPeaker outperforms other peak callers in terms of identifying known enhancers with fewer false positives. Overall, we demonstrate an optimized processing framework for STARR-seq experiments can identify putative enhancers while addressing potential confounders.

## Background

The transcription of eukaryotic genes is precisely coordinated by an interplay between *cis-*regulatory elements. For example, enhancers and promoters serve as binding platforms for transcription factors (TFs) and allow them to interact with each other via three-dimensional looping of chromatin. Their interactions are often required to initiate transcription [1,2]. Enhancers, which are often distant from the transcribed gene body itself, play critical roles in the upregulation of gene transcription. Enhancers are cell-type specific and can be epigenetically activated or silenced to modulate transcriptional dynamics over the course of development. Enhancers can be found upstream or downstream of genes, or even within introns [3–5]. They function independent of their orientation, do not necessarily regulate the closest genes, and sometimes regulate multiple genes at once [6,7]. In addition, several recent studies have demonstrated that some promoters – termed E-promoters – may act as enhancers of distal genes [8,9].

Consensus sequences (or canonical sequences) have been identified at certain protein binding sites, splice sites, and boundaries of protein-coding genes. However, there are no known consensus sequences that characterize enhancer function, making it challenging to identify enhancers based on sequence alone in an unbiased fashion. The non-coding territory occupies over 98% of the genome landscape, making the search space very broad. Moreover, the activity of enhancers depends on the physiological condition and epigenetic landscape of the cellular environment, complicating a fair assessment of enhancer function.

Previously, putative regulatory elements were computationally predicted, indirectly, by profiling DNA accessibility (using DNase-seq, FAIRE-seq, or ATAC-seq) as well as histone modifications (ChIP-seq) that are linked to regulatory functions [10–12]. More recently, researchers have developed high-throughput episomal (exogenous) reporter assays to directly measure enhancer activity across the whole genome, specifically massively parallel reporter assays (MPRA) [13,14] and self-transcribing active regulatory region sequencing (STARR-seq) [15,16]. These assays allow for quantitative assessment of enhancer activity in a high-throughput fashion.

In STARR-seq, candidate DNA fragments are cloned downstream of a reporter gene into the 3′ untranslated region (UTR). After transfecting the plasmid pool into host cells, one can measure the regulatory potential by high-throughput sequencing of the 3′ UTR of the expressed reporter gene mRNA. These exogenous reporters enable accurate and unbiased assessment of enhancer activity at the whole-genome level, independent of chromatin context. Unlike MPRA – which utilizes barcodes – STARR-seq produces self-transcribed RNA fragments that can be directly mapped onto the genome (we call this STARR-seq output hereafter). The activities of enhancers are measured by comparing the amount of RNA produced from the relative amount of genomic DNA in the STARR-seq library (we call this STARR-seq input hereafter). STARR-seq has several technical advantages over MPRA. Library construction is relatively simple because barcodes are not needed. In addition, candidate enhancers are cloned instead of synthesized, allowing the assay to test extended sequence contexts (>500 bp) for enhancer activity, which studies have shown to be critical for functional activity [17]. Importantly, STARR-seq can be scaled to the whole-genome level for unbiased scanning of functional activities. However, scaling STARR-seq to the human genome is still very challenging, primarily due to its massive size. A more complex genomic DNA library, a higher sequencing depth, and increased transfection efficiency are required to cover the whole human genome [16], which could ultimately introduce biases. Furthermore, inserting a large fragment of DNA into the 3’ UTR of the reporter gene could inadvertently introduce regulatory sequences that might affect mRNA abundance and stability, which could lead to both false positives and false negatives. MPRA is more robust in this regard because the activity of each candidate enhancer is quantified by multiple molecular barcodes associated with the fragment, making it less prone to such artifacts than STARR-seq.

The processing of STARR-seq data is somewhat similar to that of ChIP-seq, where protein-crosslinked DNA is immunoprecipitated and sequenced. A typical ChIP-seq processing pipeline identifies genomic regions over-represented by sequencing tags in a ChIP sample compared to a control sample. STARR-seq data is compatible with most ChIP-seq peak callers. Hence, previous studies on STARR-seq have largely relied on peak-calling software developed for ChIP-seq such as MACS2 [16,18,19]. However, one must be cautious using ChIP-seq peak callers, at least without re-tuning the default parameters optimized for processing TF ChIP-seq [20].

In this paper, we describe key differences in the processing of STARR-seq versus ChIP-seq data. Due to increased complexity of the genomic screening library and sequencing depth requirements, STARR-seq coverage is highly non-uniform. This leads to a lower signal-to-noise ratio than a typical ChIP-seq experiment and makes estimating the background model more challenging, which could ultimately lead to false-positive peaks. In addition, STARR-seq measures more of a continuous activity, similar to quantification in RNA-seq, than a discrete binding event. Therefore, STARR-seq peaks should be further evaluated using a notion of activity score. These differences necessitate a unique approach to processing STARR-seq data.

We propose an algorithm optimized for processing and identifying functionally active enhancers from STARR-seq data, which we call STARRPeaker. This approach statistically models the basal level of transcription, accounting for potential confounding factors, and accurately identifies reproducible enhancers. We applied our method to two whole human STARR-seq datasets and evaluated its performance against previous methods. We also compared an R package, BasicSTARRseq, developed to process peaks from the first STARR-seq data [15], which models enrichment of sequencing reads using a binomial distribution. We benchmarked our peak calls against known human enhancers. Thus, our findings support that STARRPeaker will be a useful tool for uniformly processing STARR-seq data.

## Results and Discussion

### Precise measurement of STARR-seq coverage

We binned the genome using a sliding window of length, *l*, and step size, *s*. Based on the average size of the STARR-seq library, we defined a 500 bp window length with a 100 bp step size to be the default parameter. Based on the generated genomic bins, we calculated the coverage of both STARR-seq input and output mapped to each bin. For calculating the sequence coverage, other peak callers and many visualization tools commonly use the start position of the read [15,21,22]. However, given that the average size of the fragments inserted into the STARR-seq libraries were approximately 500 bp, we expected that the read coverage using the read start position may shift the estimate of the summit of signal and dilute the enrichment. Some peak callers have used read densities of forward and reverse strands separately to overcome this issue [23,24]. To precisely measure the coverage of STARR-seq input and output, we first inferred the size of the fragment insert from paired-end reads and used the center of the fragment insert, instead of start position of the read, to calculate coverage. For inferring the size of the fragment insert, we first strictly filtered out reads that were not properly paired and chimeric. Chimeric alignments are reads that cannot be linearly aligned to a reference genome, implying a potential discrepancy between the sequenced genome and the filtered out read pairs that had a fragment insert size greater than *l*_max_ and less than reference genome and indicative of a structural variation or a PCR artifact [25]. We also filtered out read pairs that had a fragment insert size greater than *l*_max_ and less than *l*_min_. By default, we filtered out fragment insert sizes less than 200 bp and greater than 1,000 bp. After filtering out spurious read-pairs, we estimated the center of the fragment insert and counted the fragment depth for each genomic bin. To assess the benefit of using fragment-based coverage, we compared the coverage calculated using the center of fragment insert to an alternate model using the start position of the sequencing read. We found that the position of the peaks shifted up approximately 200 bp when we used the alternate model (Figure 1A, Supplementary Figure 1A). Such a shift caused by the read-based coverage could lead to the omission of TF binding sites located at the boundary. Moreover, we observed that the read-based coverage diluted the overall STARR-seq signal; as a result, peaks calculated based on the alternate model had lower fold enrichment and were less confident and broader in size (Figure 1B-D, Supplementary Figure 1B-D). Overall, the fragment-based coverage offered more concentrated and robust peak signal compared to the read-based coverage counting scheme. The benefit of using the center of the fragment is highlighted in Figure 1E, where we find more concise and precise peak with a higher fold enrichment using fragment-based coverage.

**Figure 1.**
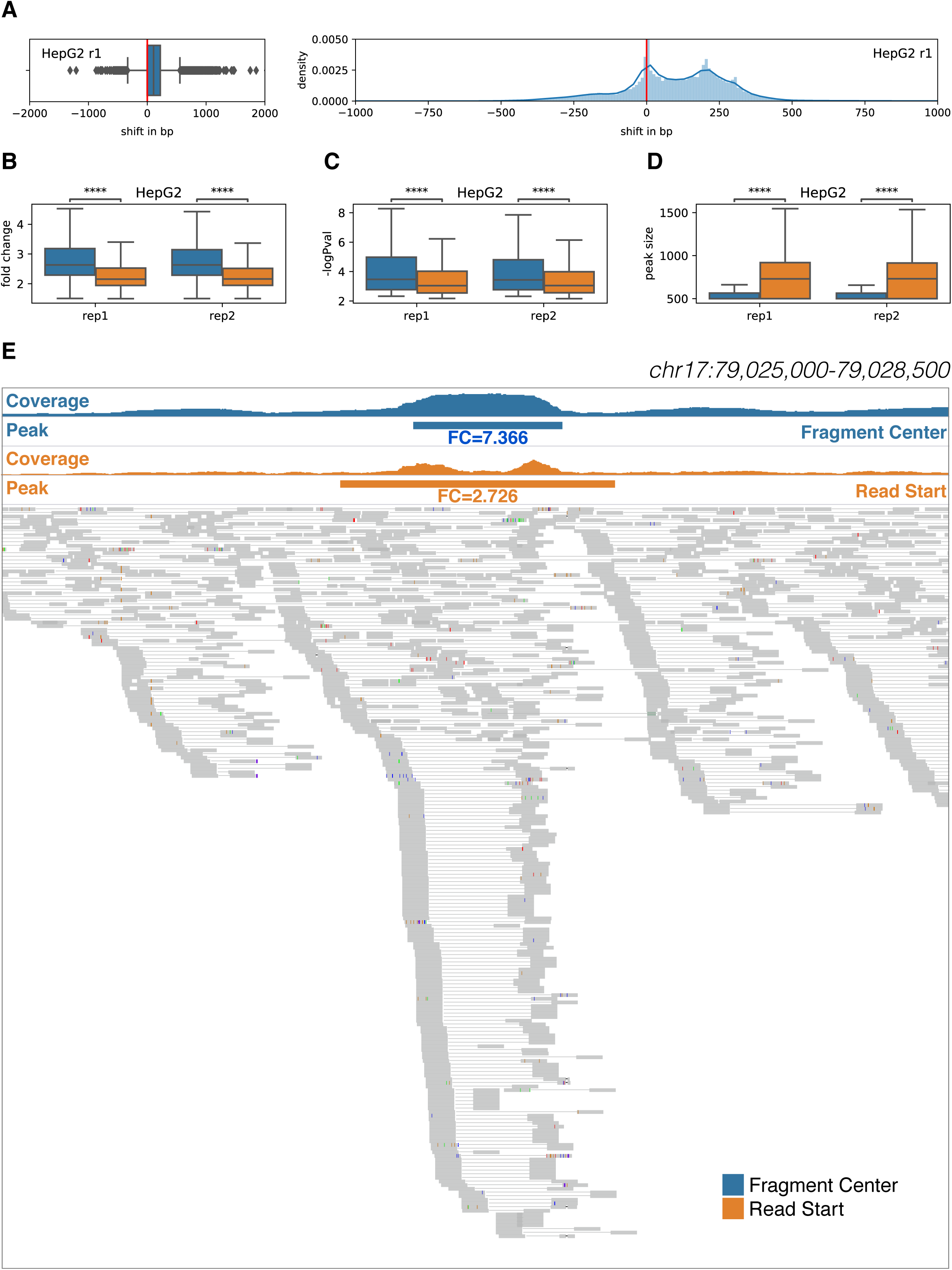
Comparison of STARR-seq output coverage calculated using the center of the fragment to using the start position of the sequencing read. **(A)** Distribution of the shift in final peak locations resulting from using two alternative coverage counting schemes in HepG2. Comparison of **(B)** overall fold enrichment level, **(C)** p-value, and **(D)** size of resulting peaks. **(E)** Example highlighting the difference between fragment-based and read-based coverage counting schemes and their resulting peak calls from HepG2 STARR-seq data. Asterisks represents statistical significance using the Mann-Whitney-Wilcoxon test two-sided with Bonferroni correction; (*) P <= 0.05, (**) P <= 0.01, (***) P <= 0.001, (****) P <= 0.0001.

### Controlling for potential systemic bias in the STARR-seq assay

To unbiasedly test for the regulatory activity, a model needs to control for potential systemic biases inherent to generating STARR-seq data. STARR-seq measures the ratio of transcribed RNA to DNA for a given test region and determines whether the test region can facilitate transcription at a higher rate than the basal level. This is based on the assumption that (1) the basal transcriptional level stays relatively constant across the genome and (2) the transcriptional rate is a reflection of the regulatory activity of the DNA insert. However, these assumptions may not always be true, and one needs to consider potential systemic biases that can interfere with the quantification of regulatory activity when analyzing the data.

We next tested whether potential sequencing biases and other covariates confounded STARR-seq readouts (Figure 2). We found that STARR-seq RNA coverage was significantly correlated with GC content (PCC 0.61; P-val 1E-299) and mappability (PCC 0.45; P-val 2.9E-148). This could be attributed to intrinsic sequencing biases in library preparation. A genome-wide reporter library is made from randomly sheared genomic DNA, but DNA fragmentation is often non-random [26]. Studies also have suggested that epigenetic mechanisms and CpG methylation may influence fragmentation [27]. Furthermore, the isolated polyadenylated RNAs are reverse transcribed and PCR amplified before sequencing, and this process can further confound the sequenced candidate fragments.

**Figure 2.**
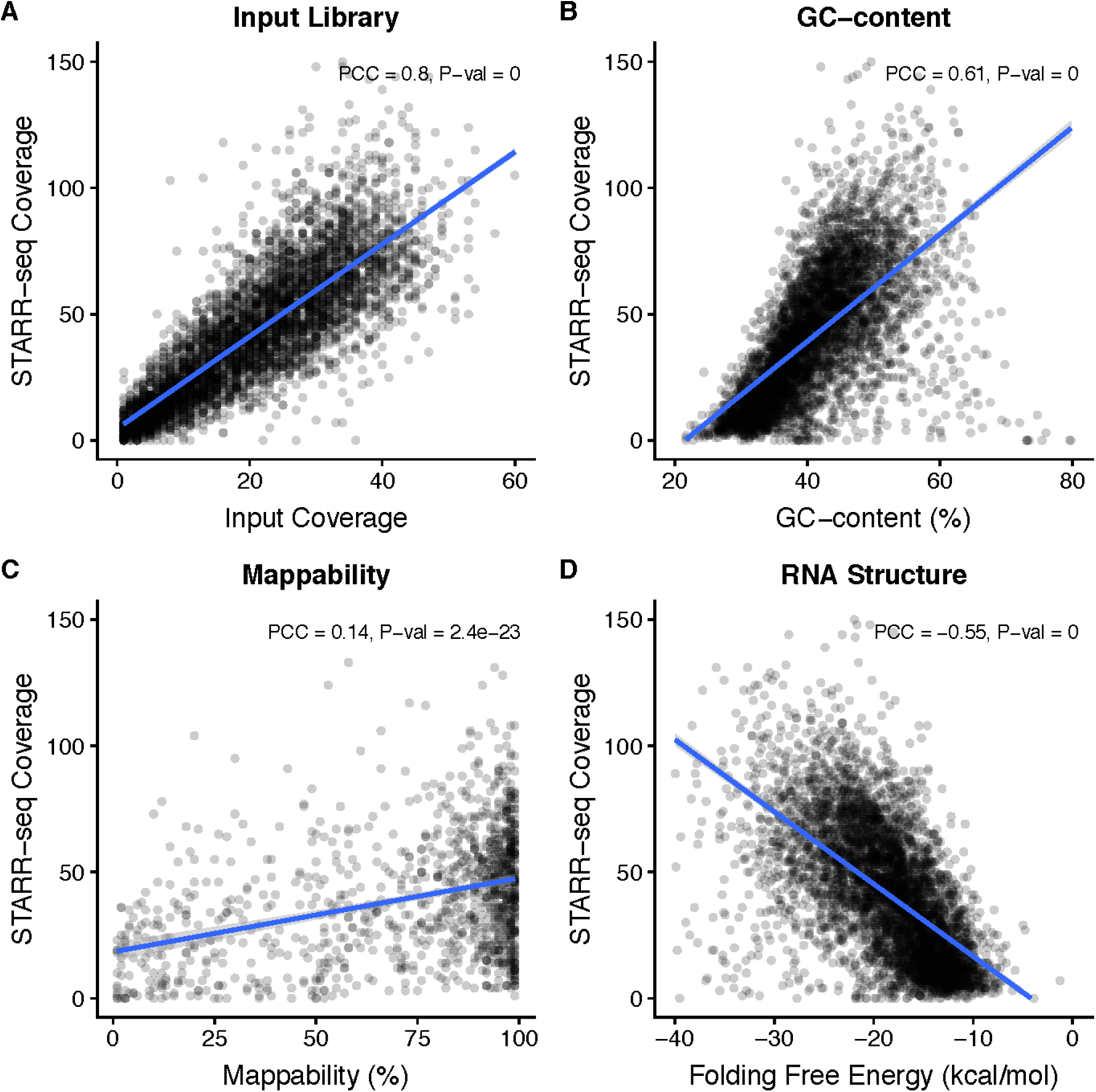
Confounding factors in the STARR-seq assay. STARR-seq output and input coverages are significantly correlated with **(A)** input coverage, **(B)** GC content, **(C)** mappability, and **(D)** RNA structure folding. PCC: Pearson Correlation Coefficient. Plots were from a sampling of 5,000 random genomic bins.

Notably, we found that STARR-seq coverage was also significantly confounded by RNA thermodynamic stability (PCC −0.55; P-val 0). Unlike ChIP-seq, where both the experiment and input controls derive from the same DNA origin, STARR-seq experiments measure the regulatory potential from the abundance of transcribed RNA, which adds a layer of complexity. For example, RNA structure and co-transcriptional folding might potentially influence the readout of STARR-seq experiments [28]. Single-stranded RNA starts to fold upon transcription and the resulting RNA structure might influence the measurement of regulatory activity. Previously, researchers suggested a potential linkage between RNA secondary structure and transcriptional regulation [29]. In addition, the resulting transcribed RNA undergoes a series of post-transcriptional regulation, and RNA stability might play a critical role. Moreover, previous reports have shown that the degradation rates – the main determinant of cellular RNA levels [30] – vary significantly across the genome and that RNA stability correlates with functionality [31,32].

Based on these findings, we built a regression-based model that accounts for various confounding variables of test sequence fragments to unbiasedly identify potential enhancer regions from STARR-seq data. Note that many of the covariates have appreciable correlation with each other. However, we did find, using stepwise forward selection, that each of them contributes substantially and independently to the model fit as assessed by Akaike information criterion (AIC) and Bayesian information criterion (BIC) (Supplementary Figure 2).

### Accurate modelling of STARR-seq using negative binomial regression

To model the fragment coverage data from STARR-seq using discrete probability distribution, we assumed that each genomic bin is independent and identically distributed, as specified in the Bernoulli trials [33]. That is, each test fragment can only map to a single fixed-length bin. Therefore, we only considered a non-overlapping subset of bins for modeling and fitting the distribution. We also excluded bins not than a minimum quantile *l*_min_, since these regions do not have sufficient power to detect covered by any genomic input or those in which the normalized input coverage was less than a minimum quantile *t*_*min*_, since these regions do not have sufficient power to detect enrichment. We selected the bin size and the minimum coverage based on the experimental design of STARR-seq. We simulated and fitted various discrete probability distributions to STARR-seq output coverage. We observed that the STARR-seq output coverage data was overdispersed and fit the best with a negative binomial distribution (Figure 3A). Moreover, a Q-Q plot of simulated coverage further demonstrated that the negative binomial model provides the best fit for the data (Figure 3B).

**Figure 3.**
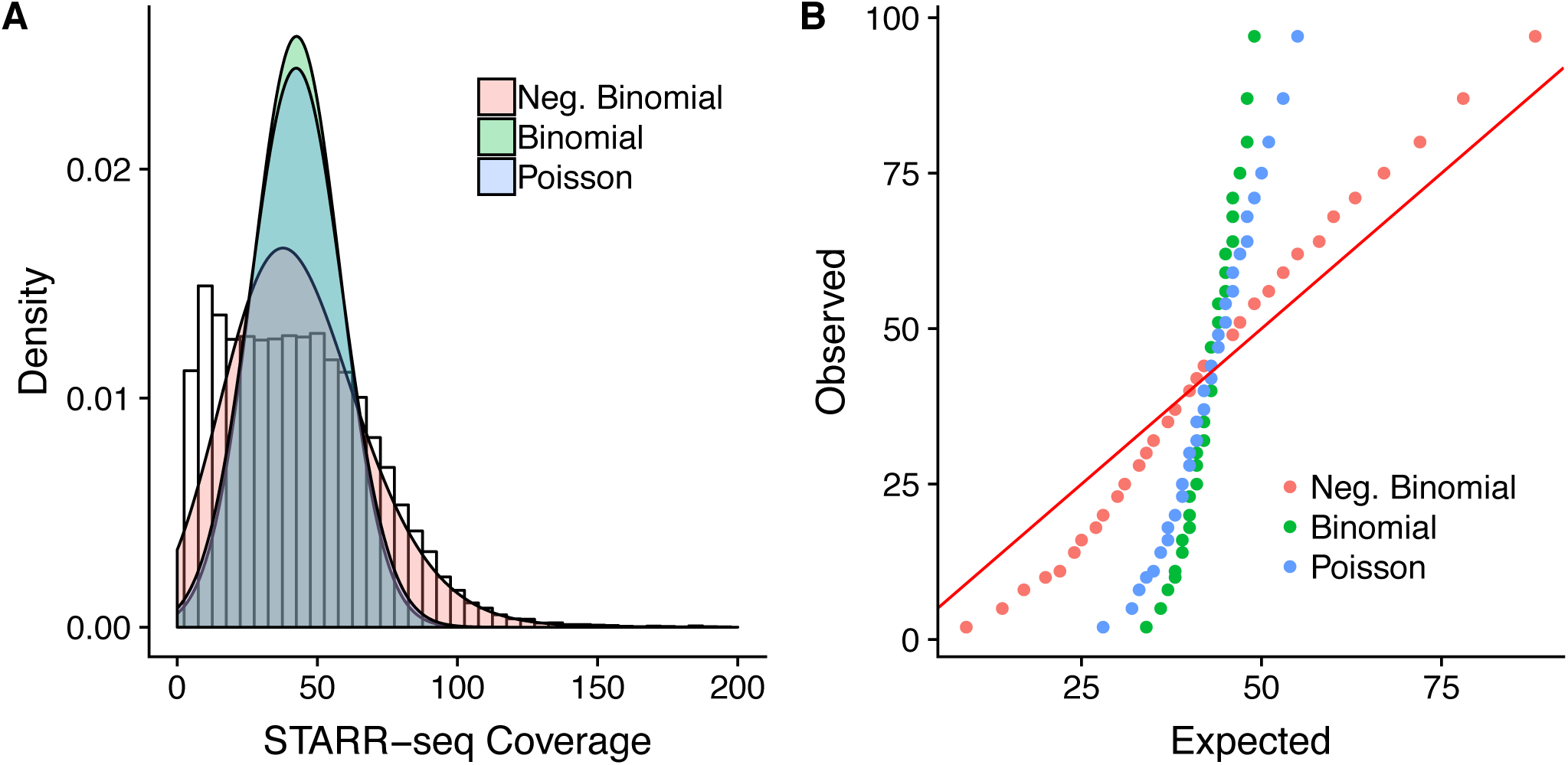
STARR-seq output coverage is fitted against simulated coverage using three distribution models; negative binomial, binomial, and Poisson. **(A)** Density histogram of simulated distribution against STARR-seq output coverage. **(B)** Q-Q plot of simulated distribution against STARR-seq output coverage. The red solid line represents where the observed count equals the expected count.

We observed a slight negative enrichment in the STARR-seq output coverage, suggesting that some candidate fragments can repress the basal transcriptional activity. However, these regions may contain sequences that can destabilize mRNAs. Therefore, additional experiments are necessary to demonstrate that STARR-seq can reliably detect silencers. In the meantime, we suggest opting for a system specifically designed for identifying silencers for this task [34].

### Peak-calling algorithm

To accurately model the ratio of STARR-seq output fragment coverage (RNA) to input fragment coverage (DNA) while controlling for potential confounding factors, we applied a negative binomial regression. The overview of our model is outlined in Figure 4. Our model starts by fitting an analytical distribution to the observed fragment coverage across fixed non-overlapping genomic bins. In doing so, we use covariates to model expected counts in the form of multiple regression. Subsequently, once a model is fitted, we evaluate the likelihood of obtaining the observed fragment counts and assign p-values using the null negative binomial distribution. In this testing phase, we use flexible genomic bins with a sliding window in order to find enrichment peaks at a higher resolution. Genomic bins with significant enrichments are selected based on their adjusted p-values using multiple testing correction. Finally, peak locations are fine-tuned to the summit of the direct fragment coverage. Note that the adjusted p-value should be regarded as the likelihood of a candidate region being an enhancer while the fold enrichment can be directly interpreted as a quantitative measure of enhancer activity.

**Figure 4.**
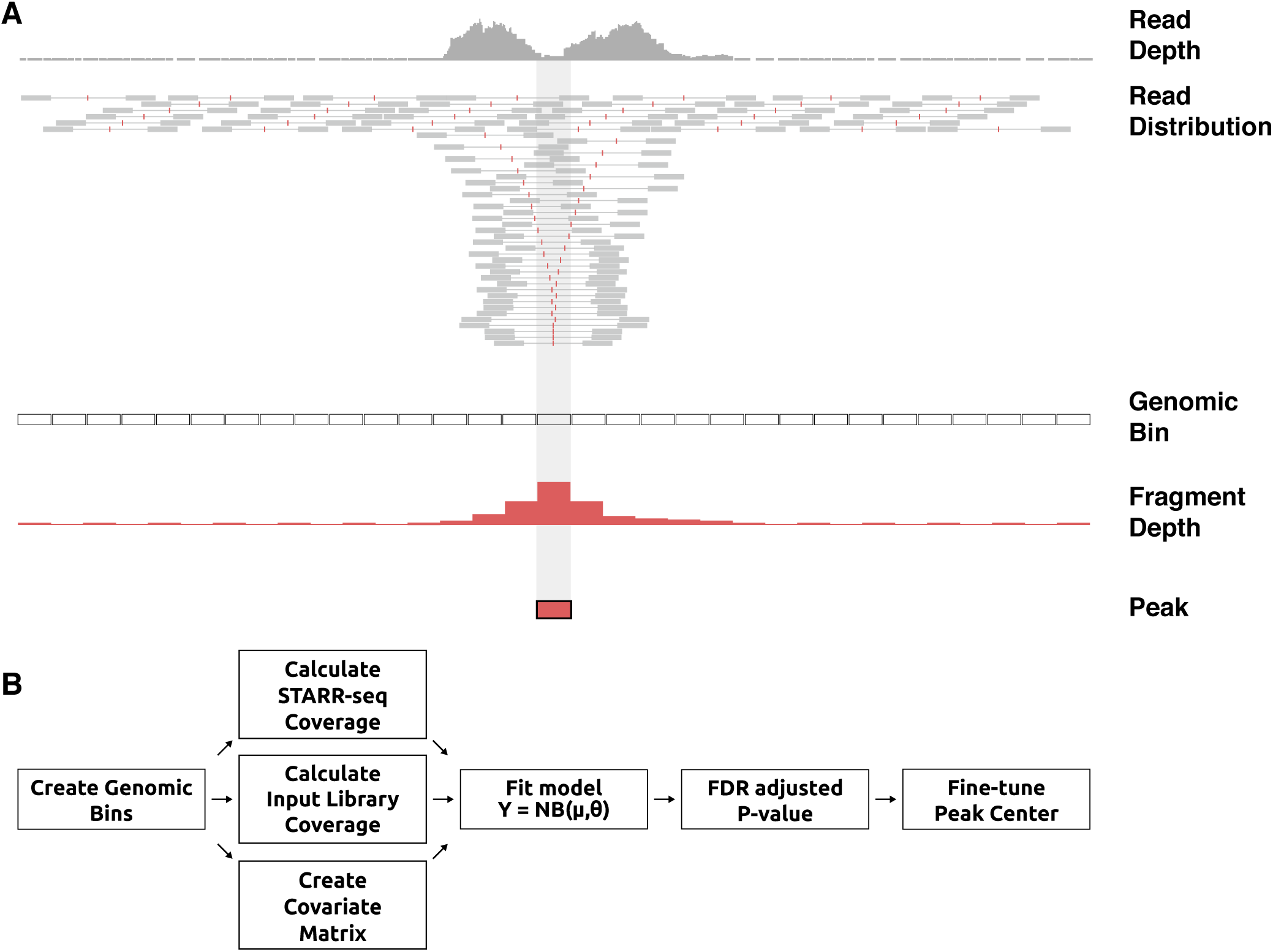
Overview of STARRPeaker peak-calling scheme. **(A)** In contrast to using read depth (grey), fragment depth (red) offers more precise and sharper STARR-seq output coverage. Fragment inserts are directly inferred from properly paired-reads. **(B)** Workflow of STARRPeaker describing how coverage is calculated for each genomic bin and modelled using a negative binomial regression model. The analysis pipeline can largely be divided into four steps: (1) Binning the genome; (2) calculating coverage and computing covariate matrix; (3) fitting the STARR-seq data to the NB regression model; and (4) peak calling, multiple hypothesis testing correction, and adjustment of the center of peaks.

Let Y be a vector of STARR-seq output (RNA) coverage, then *y*_i_ for 1 ≤ *i* ≤ *n* denotes the number of RNA fragments from a STARR-seq experiment mapped to the *i*-th bin from the total of *n* genomic bins. Let *t*_*i*_ be the number of input library fragments (DNA) mapped to the *i*-th bin. We define *X* to be the matrix of covariates, where 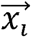 is the vector of covariates corresponding to the *i*-th bin and *x*_*ij*_ is the *j*-th covariate for the *i*-th bin.

#### Negative binomial distribution

A negative binomial distribution, which arises from a Gamma-Poisson mixture, can be parametrized as follows [35–37] (see Methods for derivation).

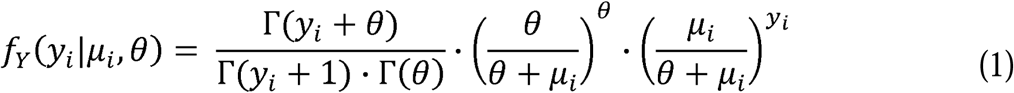

A negative binomial is a generalization of a Poisson regression that allows the variance to be different from the mean, shaped by the dispersion parameter *θ*. There are two alternative forms of parametrization for a negative binomial – NB1 and NB2 – which were first introduced by Cameron and Trivedi [36]. The difference between NB1 and NB2 is in the conditional variance of *y*_*i*_. Assuming *y*_*i*_ has mean *λ*_*i*_, the general variance function follows the form *ω* _*i*_ =*λ* _*i*_ +*αλ*_*i*_^*p*^, where *α* is a scalar parameter. NB1 uses *p*=1, whereas NB2 uses the quadratic form of variance with *p*=2. We use the most common implementation of the negative binomial, NB2, hereafter. The variance for the NB2 model is given as

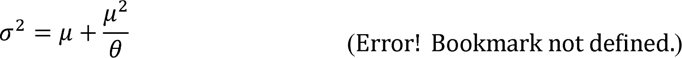

We assume that the majority of genomic bins will have a basal level of transcription, the expected fragment counts at each *i*-th bin, *E*(*y*_i_), represents the mean incidence, *μ*_*i*_, and the count of RNA fragments *Y* follows the traditional negative binomial (NB2) distribution.

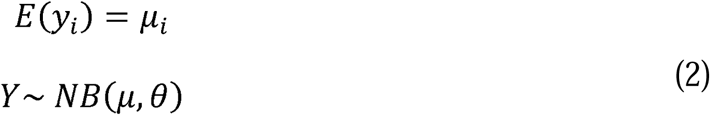

#### Negative binomial regression model

The regression term for the expected RNA fragment count can be expressed in terms a of linear combination of explanatory variables, a set of *m*covariates 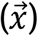. We use the input library variable *t*_*i*_ as one covariate. For simplicity, we denote *t*_*i*_ as *x*_0*i*_ hereafter.

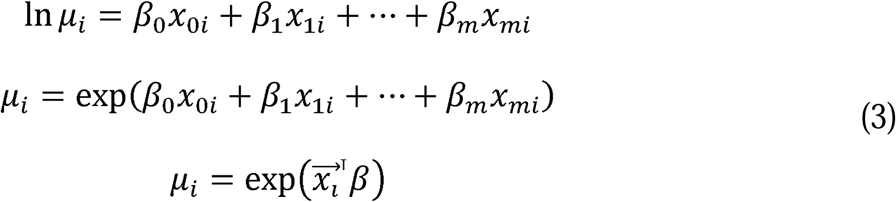

Alternatively, instead of using the input library variable *t*_*i*_ as one covariate, we can directly use it as an offset variable. Generally, a fractional observation cannot be modeled using discrete probability. However, an offset variable in a generalized linear model can be used to correct the response term to behave like a fraction. One advantage of using the input variable as an “exposure” to the RNA output coverage is that it allows us to directly model the basal transcription rate (the ratio of RNA to DNA) as a rate response variable. More details on this alternative parametrization are included in the Methods section. In our STARRPeaker model, we used four covariates; fragment coverage of input genomic libraries, GC content, mappability, and the thermodynamic stability of genomic libraries.

#### Maximum-likelihood estimation

We fit the model and estimate regression coefficients using the maximum likelihood method, where log-likelihood function is shown as follows.

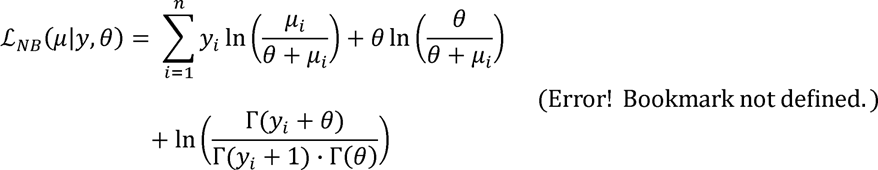

Substituting *μ*_*i*_ with the regression term, the log-likelihood function can be parametrized in terms of regression coefficients, *β*.

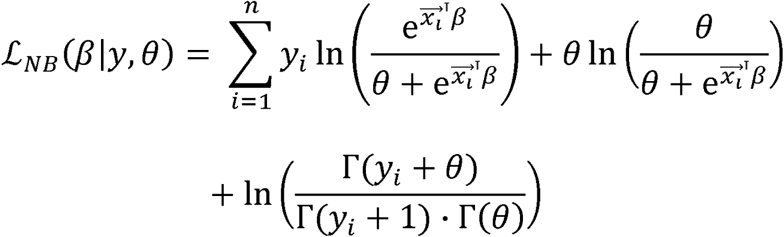

We can determine the maximum likelihood estimates of the model parameters by setting the first derivative of the log-likelihood with respect to β, the gradient, to zero, and there is no analytical solution for 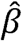. Numerically, we iteratively solve for the regression coefficients β and the dispersion parameter *θ*, alternatively, until both parameters converge.

#### Estimation of P-value

The P-value is defined as the probability of observing equal or more extreme value than the observed value at the *i*-th bin, *y*_*i*_, under the null hypothesis.

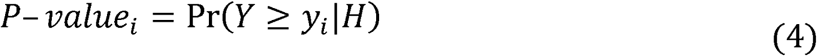

As defined earlier, we assume the random variable Y comes from a negative binomial distribution with the fitted mean at the *i*-th bin, *μ*_*i*_, as the expected value, and *θ* as the dispersion parameter. Then, we can estimate the P-value from the cumulative distribution function *CDF*, which is the sum of the probability mass function *f*_*Y*_ from 0 to *y*_*i*_ =1.

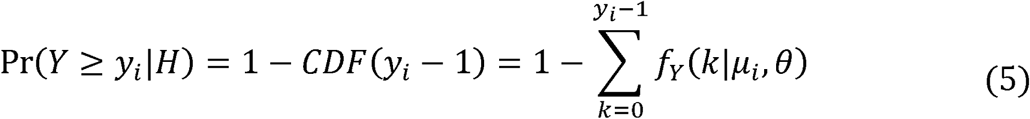

Substituting (1) gives

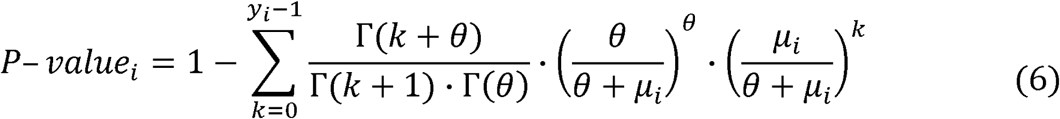

Finally, we calculate the false discovery rate using *Benjamini & Hochberg* method [38].

### Application of STARRPeaker

We applied STARRPeaker to two whole human genome STARR-seq experiments, K562 and HepG2, utilizing origin of replication (ORI)-based plasmids [39]. Based on peaks identified from these datasets, we evaluated the quality and characteristics of the identified enhancers as well as the performance of the peak caller by comparing to external enhancer resources.

#### Initial evaluation of STARRPeaker enhancers

We processed two biological replicates from each cell type independently and assessed the correlation between each pair. Overall, we observed high correlation between two replicates (PCC=0.99 for both HepG2 and K562; see Supplementary Figure 3). By intersecting peaks from two replicates, we identified 32,929 and 20,471 reproducible candidate enhancers from HepG2 and K562, respectively (Supplementary Table S1). Although the total number of peaks varied between HepG2 and K562, we observed a comparable number of peaks within the accessible region of the genome. We found 12,019 (36.34%) and 11,420 (55.57%) candidate enhancers from HepG2 and K562, respectively, within the open chromatin defined by ENCODE DNase-seq hotspots. Consistent with previous findings [39], a substantial fraction of candidate enhancers was epigenetically silenced at the chromatin level. However, as demonstrated previously using a histone deacetylase inhibitor (HDAC) [16], these poised enhancers can become functional under a more transcriptionally permissive environment. Therefore, episomal reporter assays like STARR-seq have the unique advantage of detecting potential enhancer activity independent from chromatin context. We would like to note that it is important to identify poised enhancers located in heterochromatic regions of the genome, which could become functional during developmental or pathological time courses.

#### Assessment of robustness and reproducibility of the method

A reliable peak-calling method should be able to identify peaks from suboptimal datasets. To evaluate the robustness of STARRPeaker, we used subsets of the whole-genome STARR-seq library to call peaks and compared the results. We subsampled randomly at various rates from 20 to 80% of the total dataset and compared the quality of peaks. We found that STARRPeaker was able to reliably identify the peaks using approximately 60% of the original sequencing library (Supplementary Figure 4). However, the quality of the peak calls started to deteriorate when 40% or less were used.

#### Evaluation of potential orientation bias in candidate enhancers

In general, enhancers are thought to function independent of orientation [40]. However, the fragment counts in one orientation could be skewed over the other due to orientation-specific activities, PCR, or sequencing artifacts. To test for potential orientation-based biases, we ran a binomial test on the candidate enhancers we identified. We observed a small fraction of candidate enhancers showing strand bias [3.19% for HepG2 rep1 (n=1,605); 3.76% for HepG2 rep2 (n=1,991); 7.77% for K562 rep1 (n=2,347); 5.25% for K562 rep2 (n=2,195); FDR ≤ 0.01] (Supplementary Figure 5). Less than one third of the enhancers (n=690) showed strand-specific activity in both replicates. Thus, we conclude that there is insufficient evidence to show that orientation-dependent biases are present in our STARR-seq data. Furthermore, this finding provides further support that enhancers function independent of orientation.

#### Performance comparison to other peak-calling algorithms

We evaluated the performance of STARRPeaker by comparing it to previously used methods, namely BasicSTARRseq and MACS2.

First, we qualitatively assessed the peak-calling algorithms using a simulated dataset where the ground truth exists. We created a STARR-seq dataset that consists of four spike-in controls (hybrid of DNA input library and RNA output library of known specific location). All three methods successfully identified the four control peaks with high confidence (Supplementary Figure 6). However, we noticed that BasicSTARRseq peaks were fragmented due to its limitation of fixed peak size. Moreover, the peaks were shifted toward the enrichment of sequencing reads. Furthermore, BasicSTARRseq identified a false-positive peak, and as a result, identified a total of eight regions instead of four.

Second, we quantitatively assessed the peak-calling algorithms using the whole human genome STARR-seq dataset. After uniformly calling peaks from each method using the recommended default settings, we evaluated the quality of the candidate enhancers identified. We found that both BasicSTARRseq and MACS2 called significantly more peaks (4 to 20-fold higher) than STARRPeaker (Supplementary Table S4). While it is uncertain how many true enhancers were present in each sample, we had to ensure that we made a fair comparison across different methods due to the tradeoff between sensitivity and specificity. An increase in sensitivity is generally achieved at the expense of a decrease in specificity, as described in receiver operating characteristic curves. In our context, a method having higher specificity suffers from having less overlap with open chromatin and previously identified enhancers from other assays. Therefore, we used a uniform P-value threshold of 0.001 and subsampled the peaks before the comparison. After uniformly processing the dataset using each method, we measured the level of epigenetic profile enrichment around the peaks. We observed higher enrichment of DNase-hypersensitive sites, as well as more distinct double-peak patterns of H3K27ac and H3K4me1, using STARRPeaker compared to BasicSTARRseq or MACS2 (Figure 5, Supplementary Figure 7). Furthermore, STARRPeaker peaks had significantly higher enrichment of TF binding events (based on the number of TF ChIP-seq binding sites) compared to the peaks identified using other methods.

**Figure 5.**
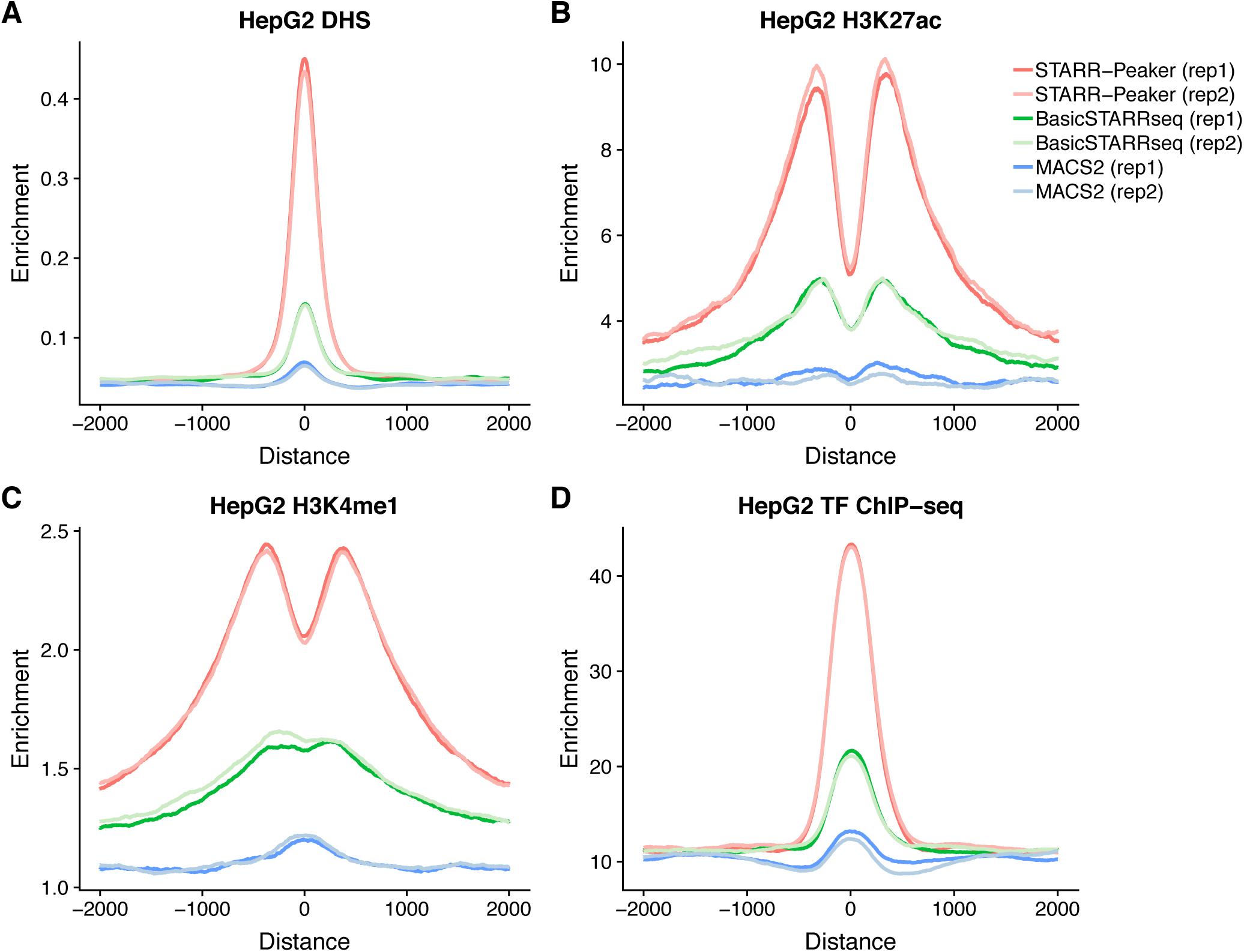
Enrichment of epigenetic signals around peaks in HepG2. All peaks were centered at the summit, uniformly thresholded using P-value < 0.001, and 10,000 peaks were randomly selected. Aggregated read depth at 2,000 bp upstream and downstream were plotted for **(A)** DNase I hypersensitive sites (DHS), **(B)** H3K27ac, **(C)** H3K4me1, and **(D)** aggregated TF ChIP-seq profile. For DNase-seq, enrichment indicates unique read depth. For histone ChIP-seq, enrichment indicates fold change over control. For TF ChIP-seq aggregate, enrichment indicates the number of TFs binding.

#### Comparison to previously characterized enhancers

First, we compared the peaks identified by STARRPeaker to previously characterized enhancers from HepG2 or K562 cell lines by CAGE [41], MPRA [17,42], and STARR-seq [19] (Figure 6, Supplementary Table S2). Overall, we observed a higher fraction of STARRPeaker peaks overlapping with external datasets compare to other methods. Moreover, we found higher overlaps when peaks from both replicates were merged, due to fewer but more precise candidate enhancers from merging replicates. However, we noticed reduced agreement across different types of enhancer assays. Low overlap between assays may arise from different formats or layouts of reporter plasmids, such as differing enhancer cloning sites or promoters, or differences in the complexity of the screening library. Furthermore, CAGE is an entirely different assay from episomal reporter assays like MPRA and STARR-seq, with enhancers defined based on bidirectional transcripts originating from an eRNA.

**Figure 6.**
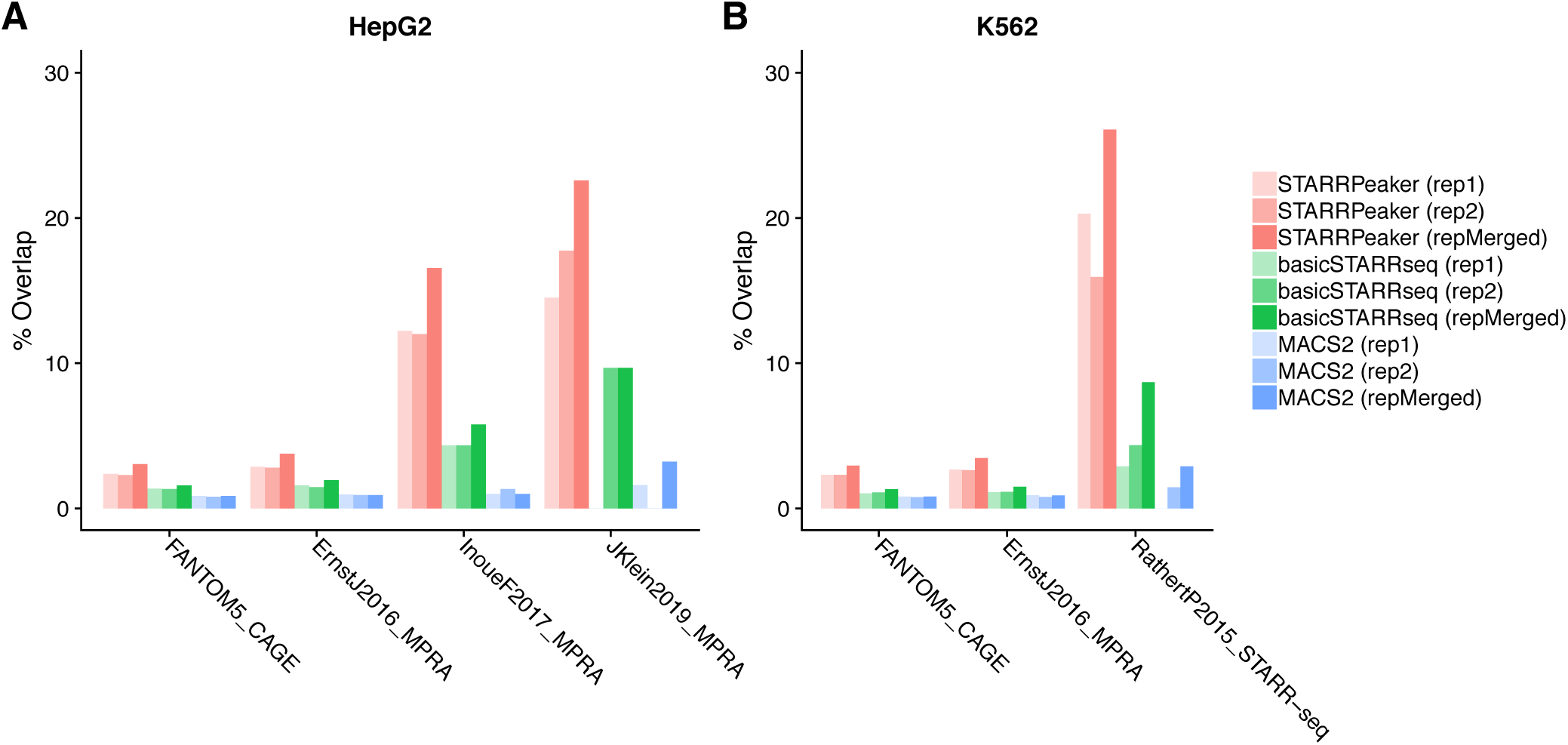
Comparison of peaks using an external dataset for **(A)** HepG2 or **(B)** K562 cell lines. Peaks identified from STARRPeaker as well as BasicSTARRseq and MACS2 were compared against a published dataset. For a fair comparison, all peaks were centered at the summit, uniformly thresholded using P-value < 0.001, and 20,000 peaks were randomly drawn from peaks identified by each peak caller using the recommended settings. The fraction of overlap was computed for each replicate. We considered it an overlap when at least 50% of peaks intersected each other.

Second, we examined the nine distal enhancers from the GATA1 and MYC loci characterized in-depth by CRISPRi tiling screen (Supplementary Figure 8). We found that STARRPeaker accurately called peaks for 6 of 9 enhancers from both replicates. For the remaining three regions, we observed insufficient enrichment of STARR-seq output and, therefore, we concluded that this is not a shortcoming of the peak caller.

#### Application to external STARR-seq datasets

To ensure that STARRPeaker can be generally applied to different variants of STARR-seq assays, we tested STARRPeaker on previously published STARR-seq datasets. First, we applied STARRPeaker to the whole-genome ORI-STARR-seq dataset on HeLa-S3 [39] and assessed the quality of the peaks identified. Consistent with the previous claim that IFN-I signaling may induce false-positive enhancers, we identified more peaks in untreated HeLa-S3 samples (n=28,381) compared to inhibitor-treated samples (n=16,150). Furthermore, peaks from untreated samples had lower enrichment of chromatin accessibility (DNase-seq) than those from inhibitor-treated samples, supporting that TBK1/IKK/PKR inhibition reduces false-positive enhancer signals related to IFN-I signaling (Supplementary Figure 9A). Moreover, STARRPeaker covered 77.5% (n=7,451) of published peaks, which were called using BasicSTARRseq and then further shortlisted using a stringent threshold (P-value 1E-5 with corrected enrichment ≥ 4). Furthermore, STARRPeaker found 6,540 additional peaks from a HeLa-S3 sample that was highly enriched with chromatin accessibility signals (Supplementary Figure 9B). Second, we tested if STARRPeaker can be reliably applied to captured STARR-seq datasets (Cap-STARR-seq). We applied STARRPeaker to a previously characterized GM12878 STARR-seq dataset based on an ATAC-seq-capture technique called HiDRA [43] and compared its performance with published results. The HiDRA dataset was reported to have ∼65,000 regions with enhancer function. In the STARRPeaker run, we identified only 20,852 regions with significant enhancer activities from the five replicates they produced. Approximately 73.6% of peaks overlapped with the published results (n=15,347). While it is debatable to claim that one method is superior to the other, this result demonstrates that STARRPeaker can be reliably used against the Cap-STARR-seq dataset.

Third, we further evaluated the performance of the peak-calling methods by applying STARRPeaker and two other peak-calling methods to another published Cap-STARR-seq dataset [19]. The dataset covers approximately 91% of the surrounding 3 Mb of the MYC locus. Consistent with the earlier analysis, we observed that STARRPeaker is highly specific and identifies fewer candidate enhancers (n=26) compared to the other methods (BasicSTARRseq n=223; MACS2 n=136). Furthermore, a four-way comparison (STARRPeaker, BasicSTARRseq, MACS2, and published peaks) showed that all of the STARRPeaker peaks overlapped with peaks from other methods but not the other way around (Supplementary Figure 10). These results indicate that STARRPeaker is more robust and reliable at identifying reproducible candidate enhancers from various STARR-seq datasets than previous methods.

## Conclusions

In summary, we developed a reliable peak-calling analysis pipeline named STARRPeaker that is optimized for large-scale STARR-seq experiments. To illustrate the utility of our method, we applied it to two whole human genome STARR-seq datasets from K562 and HepG2 cell lines, utilizing ORI-based plasmids. STARRPeaker has several key improvements over previous approaches including (1) precise and efficient calculation of fragment coverage; (2) accurate modeling of the basal transcription rate using negative binomial regression; and (3) accounting for potential confounding factors, such as GC content, mappability, and the thermodynamic stability of genomic libraries. We demonstrate the superiority of our method over previously used peak callers, supported by strong enrichment of epigenetic marks relevant to enhancers and overlap with previously known enhancers.

To fully understand how noncoding regulatory elements can modulate transcriptional programs in human, STARR-seq active regions must be further characterized and validated within different cellular contexts. For example, recent applications of CRISPR-dCas9 to genome editing have allowed researchers to epigenetically perturb and test these elements in their native genomic context [44,45]. The next step for CRISPR-based functional screens is to overcome the current limitation of small scale by leveraging barcodes and single-cell sequencing technology [46]. In the meantime, we envision that the STARRPeaker framework could be utilized to detect and quantify enhancers at the whole-genome level, thereby aiding in prioritizing candidate regions in an unbiased fashion to maximize functional characterization efforts.

## Methods

### Cell culture

We cultured K562 cells (ATCC) in IMDM (Gibco #12440) supplemented with 10% fetal bovine serum (FBS) and 1% pen/strep and maintained in a humidified chamber at 37°C with 5% CO_2_. We cultured HepG2 cells (ATCC) in EMEM (ATCC #30-2003) supplemented with 10% FBS and 1% pen/strep, maintained in a humidified chamber at 37°C with 5% CO_2_.

### Generating an ORI-STARR-seq input plasmid library

We sonicated human male genomic DNA (Promega #G1471) using a Covaris S220 sonicator (duty factor – 5%; cycle per burst – 200; 40 sec) and ran it on a 0.8% agarose gel to size-select 500 bp fragments. After gel purification using a MinElute Gel Extraction kit (Qiagen), we end-repaired, ligated custom adaptors, and PCR-amplified DNA fragments using Q5 Hot Start High-Fidelity DNA polymerase (NEB) (98°C for 30 sec; 10 cycles of 98°C for 10 sec, 65°C for 30 sec, and 72°C for 30 sec; 72°C for 2 min) to add homology arms for Gibson assembly cloning.

We used AgeI-HF (NEB) and SalI-HF (NEB) to linearize the hSTARR-seq_ORI plasmid (gift from Alexander Stark; Addgene plasmid #99296) and cloned the PCR products into the vector using Gibson Assembly Master Mix (NEB); we set up 60 replicate reactions to maintain complexity. We purified the assembly reactions using SPRI beads (Beckman Coulter), dialyzed them using Slide-A-Lyzer MINI dialysis devices (ThermoScientific), and concentrated them using an Amicon Ultra-0.5 device (Amicon). We transformed the reaction into MegaX DH10BTM T1 electrocompetent cells (Thermo Fisher Scientific) (with 25 replicate transformations to maintain complexity) and let them grow in 12.5L LB-Amp medium until they reached an optical density of ∼1.0. We extracted the plasmids using a Plasmid Gigaprep Kit (Qiagen) and dialyzed the plasmid prep using Slide-A-Lyzer MINI dialysis devices before electroporation.

### Electroporation-mediated transfection of ORI-STARR-seq input plasmid library into K562 and HepG2 cell lines

We electroporated the ORI-STARR-seq library using an AgilePulse Max (Harvard Apparatus) and generated two biological replicates for each cell line. For K562 cells, we electroporated 5.6 mg of input plasmid library into 700 million cells per biological replicate by delivering three 500 V pulses (1 ms duration with a 20 ms interval). For HepG2 cells, we electroporated 8 mg of input plasmid library into one billion cells in one replicate, and 5.6 mg into 700 million cells in another replicate by delivering three 300 V pulses (5 ms duration with a 20 ms interval).

### Generation of an Illumina sequencing library

#### Output RNA library

We harvested cells 24 hr after electroporation, and extracted total RNA using an RNeasy Maxi kit (Qiagen). We further isolated polyA-plus mRNA using Dynabeads® Oligo (dT) kit (ThermoFisher Scientific), treated it with TURBO DNase (Invitrogen), and purified the reaction using an RNeasy MinElute Kit (Qiagen). We synthesized cDNA using SuperScript III (ThermoFisher Scientific) with a custom primer that specifically recognizes mRNAs that had been transcribed from the ORI-STARR-seq library. After reverse transcription, we treated the reactions with a cocktail of RNase A and RNase T1 (ThermoFisher Scientific). We split cDNA samples into 160 replicate sub-reactions, and PCR-amplified each sub-reaction with a primer with a unique index (helping to identify PCR duplicates) using Q5 Hot Start High-Fidelity DNA polymerase (NEB) with the following program: 98°C for 30 s; cycles of 98°C for 10 s, 65°C for 30 s, 72°C for 30 s (until they reached mid-log amplification phase; we cycled 18 cycles for K562 Rep.1; 16 cycles for K562 Rep. 2; 18 cycles for HepG2 Rep. 1; and 15 cycles for HepG2 Rep2); 72°C for 2 min). After PCR, we re-combined all sub-reactions into one and purified it with Agencourt Beads. We generated 100 bp paired-end reads for each biological replicate on an Illumina Hiseq4000 at the University of Chicago Genome Facility.

#### Input DNA library

We PCR-amplified a total of 200 ng of input plasmid library (in 16 replicate reactions) using Q5 Hot Start High-Fidelity DNA polymerase (NEB) with the following program: 98°C for 30 s; 4 cycles of 98°C for 10 s, 65°C for 30 s, and 72°C for 20 s; 8 cycles of 98°C for 10 s and 72°C for 50 s; 72°C for 2 min). After PCR, we combined all products into one and purified it with Agencourt Beads. We generated 100 bp paired-end reads on an Illumina Hiseq4000 at the University of Chicago Genome Facility.

### Sequencing and preprocessing

For each of 160 replicates, paired-end sequencing reads were aligned to the human reference genome GRCh38 downloaded from the ENCODE portal (ENCSR425FOI) using BWA-mem (v0.7.17). Alignments were filtered against unmapped, secondary alignments, mapping quality score less than 30, and PCR duplicates using SAMtools (v1.9) and Picard (v2.9.0). All of the replicates were pooled and sorted for downstream analysis.

### Negative binomial distribution

A negative binomial distribution, which arises from Gamma-Poisson mixture, can be parametrized for y>=0 as follows.

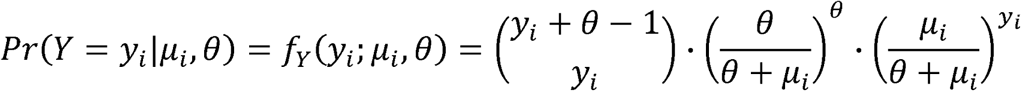

Where

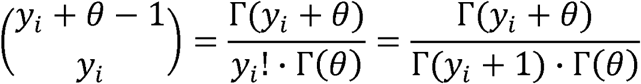

Substituting gives:

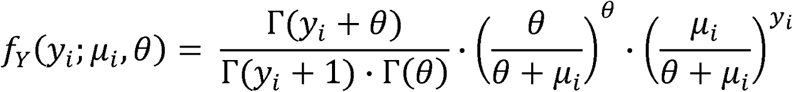

Rearranging gives:

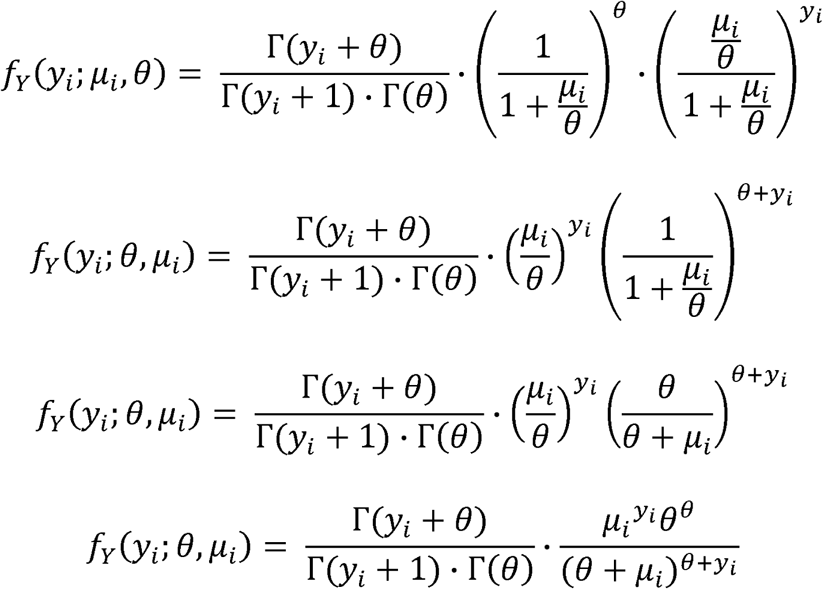

### Alternative parametrization of negative binomial regression using a rate model

Alternative parametrization allows STARR-seq data to be modelled as a rate model. In contrast to using input coverage as one of the covariates, we can consider it as “exposure” to output coverage. This “trick” allows us to directly model the basal transcription rate (the ratio of RNA to DNA) as a rate response variable. We defined the transcription rate (RNA to DNA ratio) as a new variable, π_*i*_.

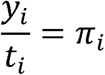

If we assume the majority of genomic bins will have the basal transcription rate, we can model the transcription rate at each *i*-th bin following the traditional negative binomial (NB2) distribution.

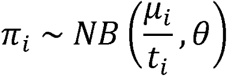

The expected basal transcription, *E (π*_*i*_), becomes the mean incidence rate of *y*_*i*_ per unit of exposure, *t*_*i*_.

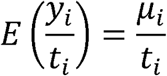

By normalizing *μ* _*i*_ by *t*_*i*_, we are modeling a rate instead of a discrete count using the negative binomial distribution. The regression term for the expected transcription rate can be expressed in terms of a linear combination of explanatory variables, *j* covariates 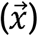.

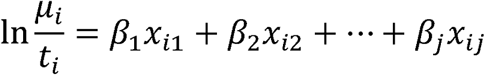

Rearranging in terms of the expected value of *y*, or *μ*, gives

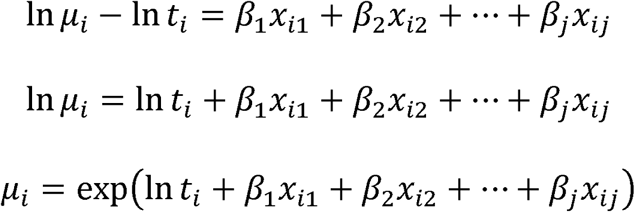

The natural log of *t*_*i*_ on the RHS ensures *μ*_*i*_ is normalized in the model, acting as an offset variable. In STARRPeaker software, we allow users to optionally choose this alternative rate model (implemented as “mode 2”) instead of the default covariate model described in the main text. This alternate model is useful if constant basal transcription is expected throughout the genome or if covariates are available for directly modelling the basal transcription rate *π*.

### BasicSTARRseq

We used BasicSTARRseq R package version 1.10.0 downloaded from Bioconductor (https://bioconductor.org/packages/release/bioc/html/BasicSTARRseq.html). We used default setting as described in the software manual, except for disabling deduplication (minQuantile = 0.9, peakWidth = 500, maxPval = 0.001, deduplicate = FALSE, model = 1), to call peaks.

### MACS2

We used MACS2 version 2.1.1 [23] at the recommended default setting, except for allowing duplicates in read (--keep-dup all), since our STARR-seq dataset was multiplexed. We called peaks with an FDR cutoff of 0.01, as recommended by the author of the software.

### Calculating folding free energy

We used the LinearFold [47] algorithm to estimate the folding energy of each genomic bin iteratively across the whole genome. Specifically, we used the Vienna RNAfold thermodynamic model [48] with parameters from Mathews et al. 2004 [49]. We implemented a parallel processing scheme to leverage multicore processors to expedite the calculation of folding free energy.

## Declarations

### Availability of data and source codes

We implemented the method described in this article as a Python software package called STARRPeaker. The software package can be downloaded, installed, and readily used to call peaks from any STARR-seq dataset. The STARRPeaker package, as well as source code and documentation, is freely available at: http://github.com/gersteinlab/starrpeaker. All raw data used in the analysis as well as derived resources are available to download from the ENCODE portal (https://www.encodeproject.org/) with accession code ENCSR135NXN for HepG2 and ENCSR858MPS for K562. DNase-seq and ChIP-seq data used for the analysis is also publicly available from the ENCODE portal. The specific accession codes used for the analysis are listed in Supplementary Table S3. GC content was downloaded from the UCSC Genome Browser (http://hgdownload.cse.ucsc.edu/gbdb/hg38/bbi/gc5BaseBw/), and the mappability track was created using gem-library software [50] with a k-mer size of 100 bp and the reference human genome build hg38.

### Author Contributions

D.L., M.S., K.W., and M.G. conceived the project. D.L. and M.G. drafted the manuscript. D.L. developed the STARRPeaker software package. M.S., J.M., M.W., D.F., Y.K., and L.M. performed experimental works. M.W. and Y.K. performed experimental validations. D.L., J.Z., and J.L. performed the downstream analyses. M.G. and K.W. provided funding and supervised the project.

## Acknowledgements

We thank Jinrui Xu and Joel Rozowsky for thoughtful discussion about ChIP-seq processing, Michael Rutenberg Schoenberg and Zhen Chen for thoughtful discussion about RNA-folding biology, and all other members of the Gerstein and White laboratories for advice and critical feedback on the manuscript.

## Supplementary Tables

**Table S1** contains significant peaks called by STARRPeaker.

**Table S2** contains overlap of various peak callers (STARRPeaker, BasicSTARRseq, and MACS2) to published enhancers identified using other types of enhancer assays.

**Table S3** contains a list of data sources and accession numbers used for the analysis.

**Table S4** compares peaks identified by various peak callers (STARRPeaker, BasicSTARRseq, and MACS2).

## Supplementary Figures

**Supplementary Figure 1.**
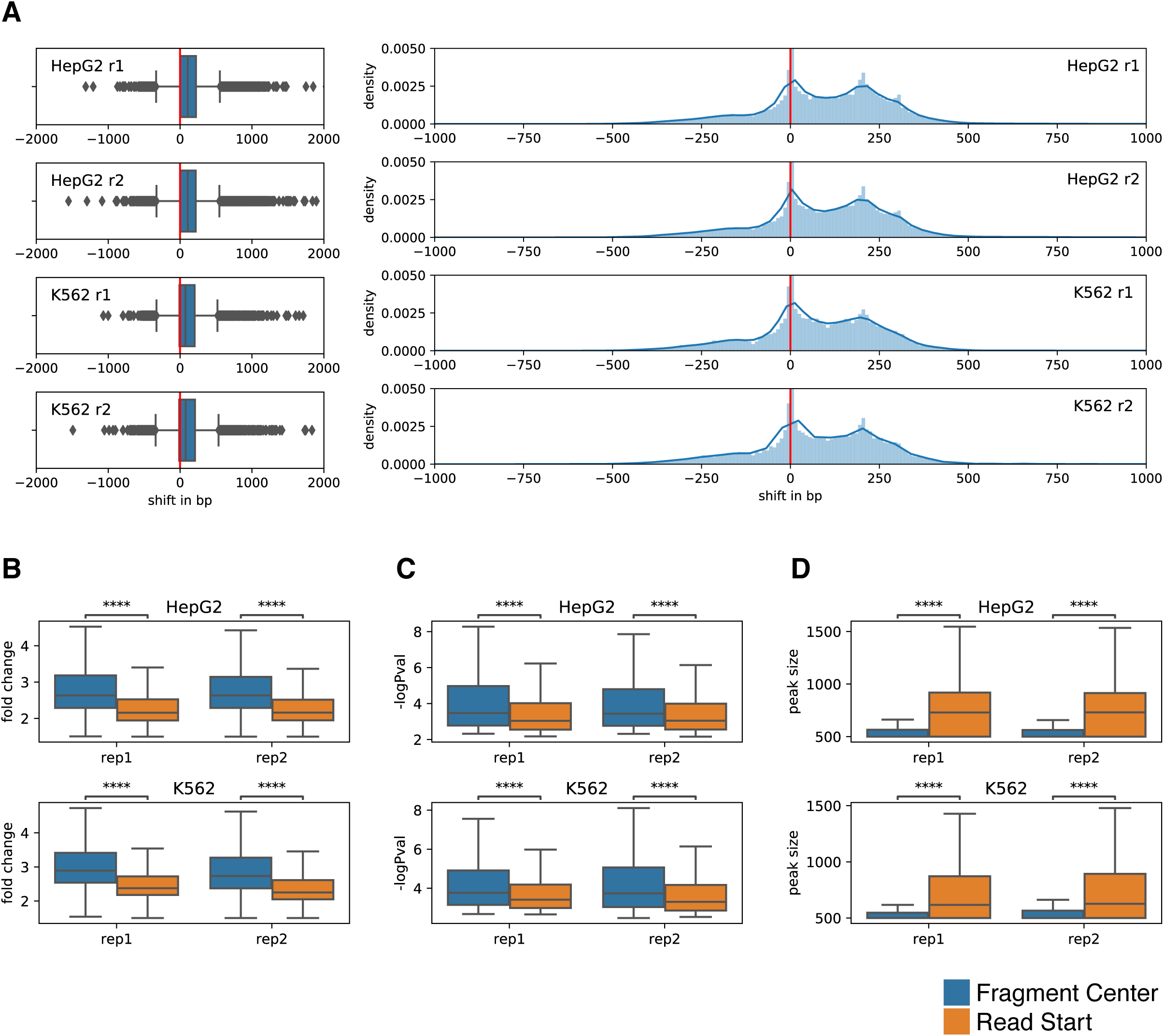
Comparison of STARR-seq output coverage calculated using the center of the fragment to using the start position of the sequencing read. **(A)** Distribution of shift in final peak locations resulting from using two alternative coverage counting schemes in HepG2. Comparison of **(B)** overall fold enrichment level, **(C)** p-value, and **(D)** size of resulting peaks.

**Supplementary Figure 2.**
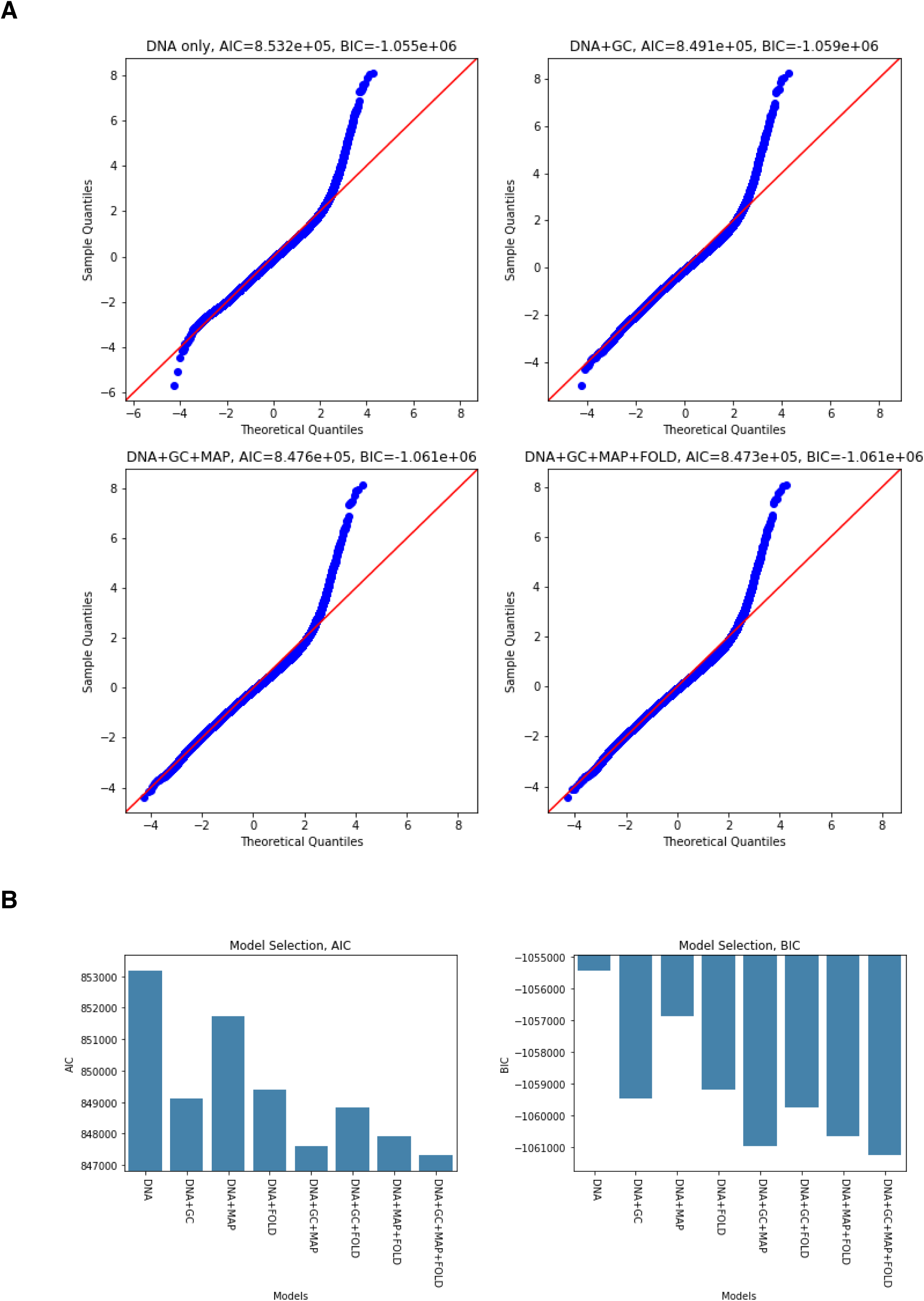
Contribution of covariates and model selection. **(A)** Q-Q plots of various models with different sets of covariates showing the goodness of fit. **(B)** Both AIC and BIC measure relative qualities of statistical models considering the trade-off between the goodness of fit and the simplicity of the model. AIC: Akaike information criterion; BIC: Bayesian information criterion.

**Supplementary Figure 3.**
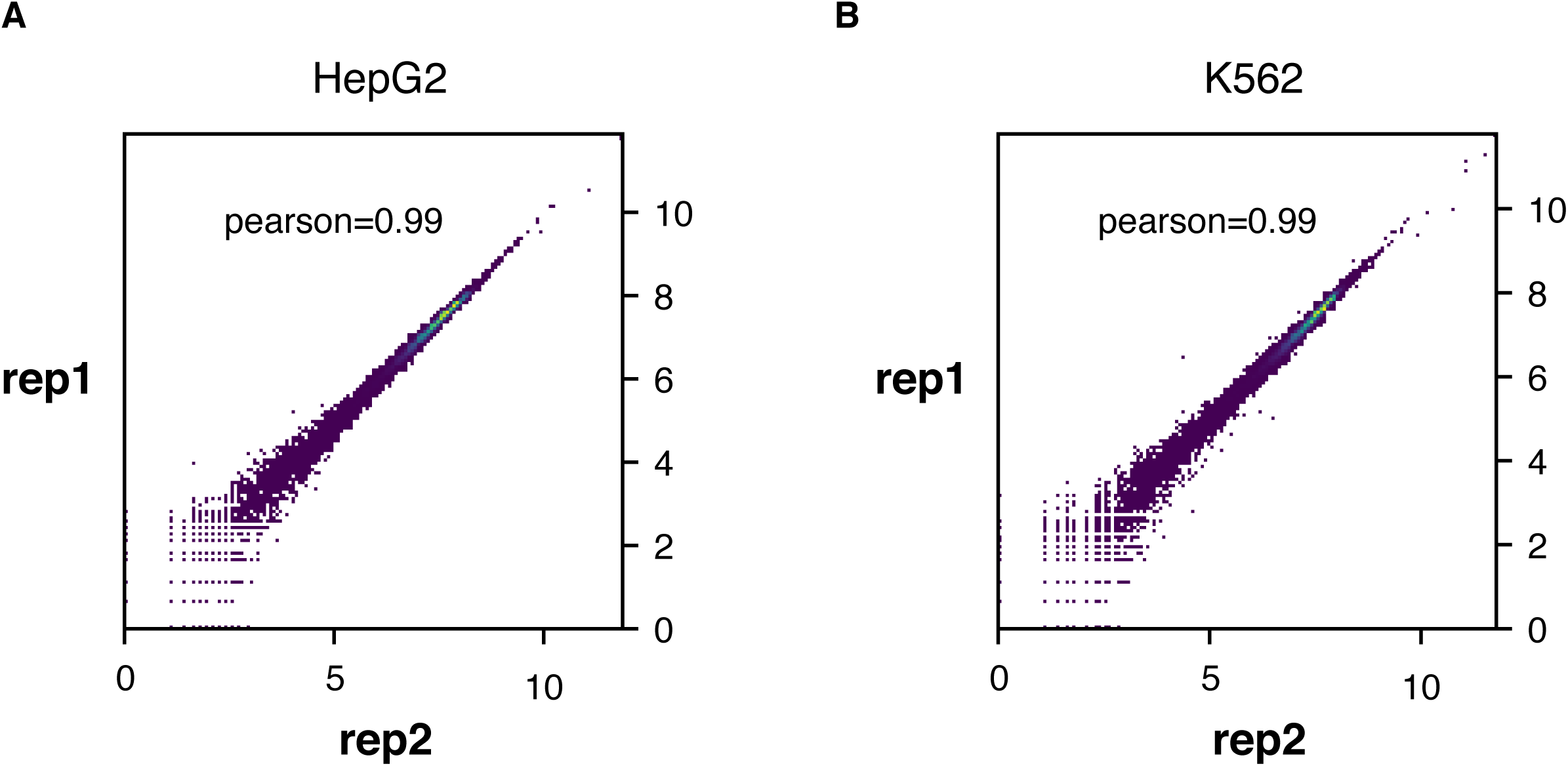
Correlation between replicates for **(A)** HepG2 or **(B)** K562 cell lines.

**Supplementary Figure 4.**
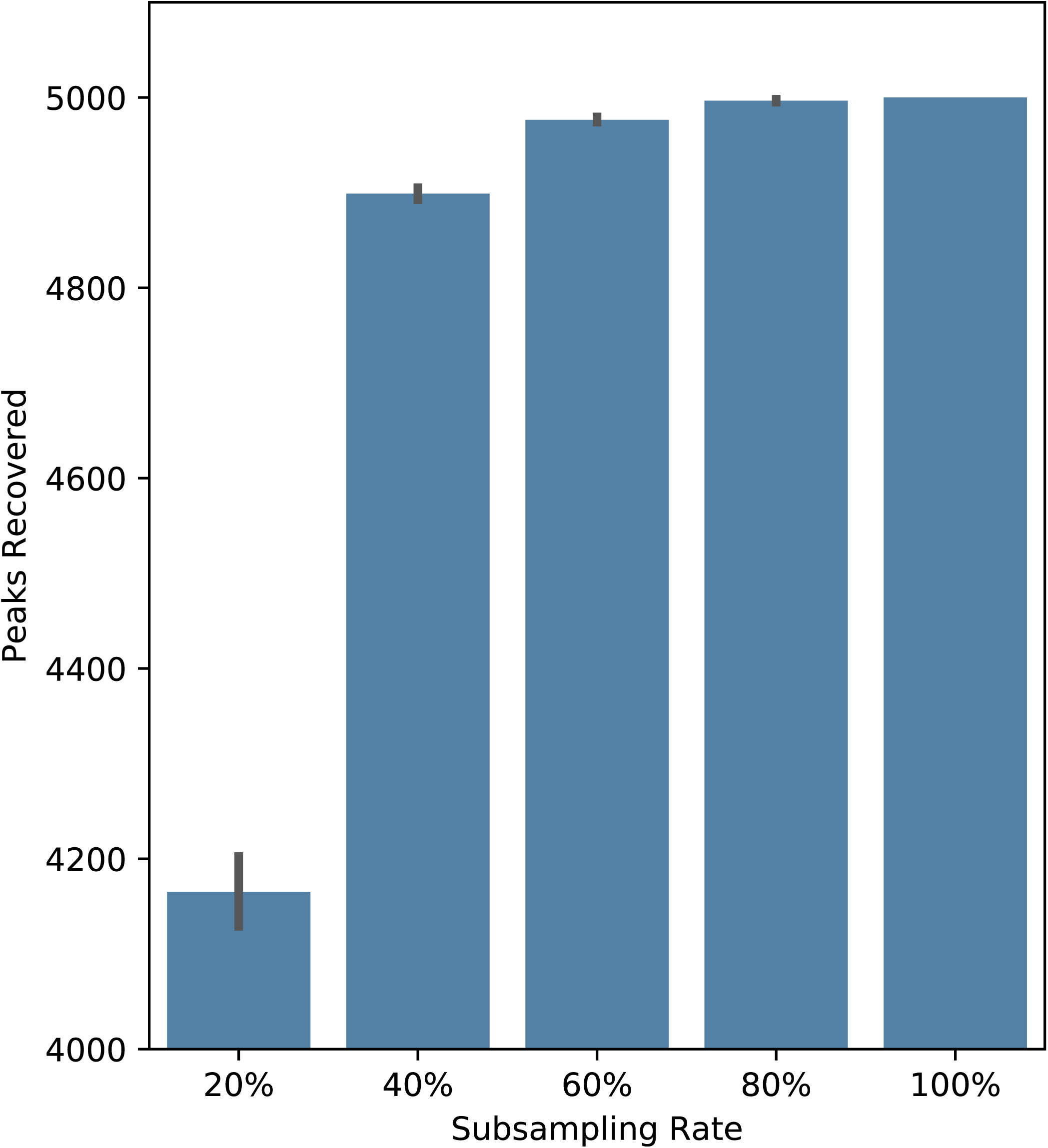
Comparison of peaks called from subsamples of the original STARR-seq library, highlighting the robustness of STARRPeaker.

**Supplementary Figure 5.**
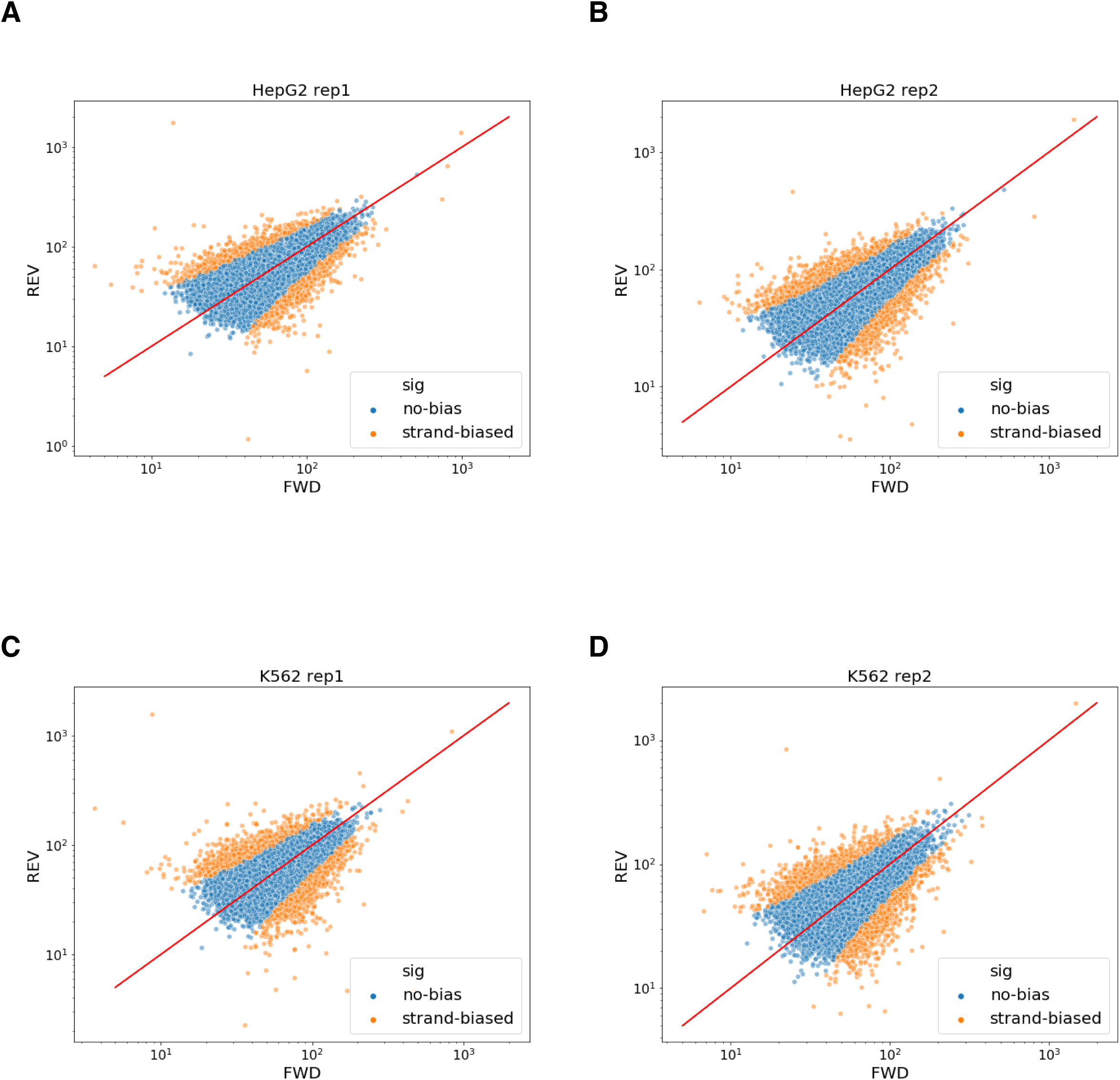
Orientation biases analysis for **(A-B)** HepG2 or **(C-D)** K562 cell lines. The ratio between forward and reverse stranded fragments was tested for statistical significance using a binomial test. Orange dots represent peaks with significant strand bias (FDR q-value < 0.01).

**Supplementary Figure 6.**
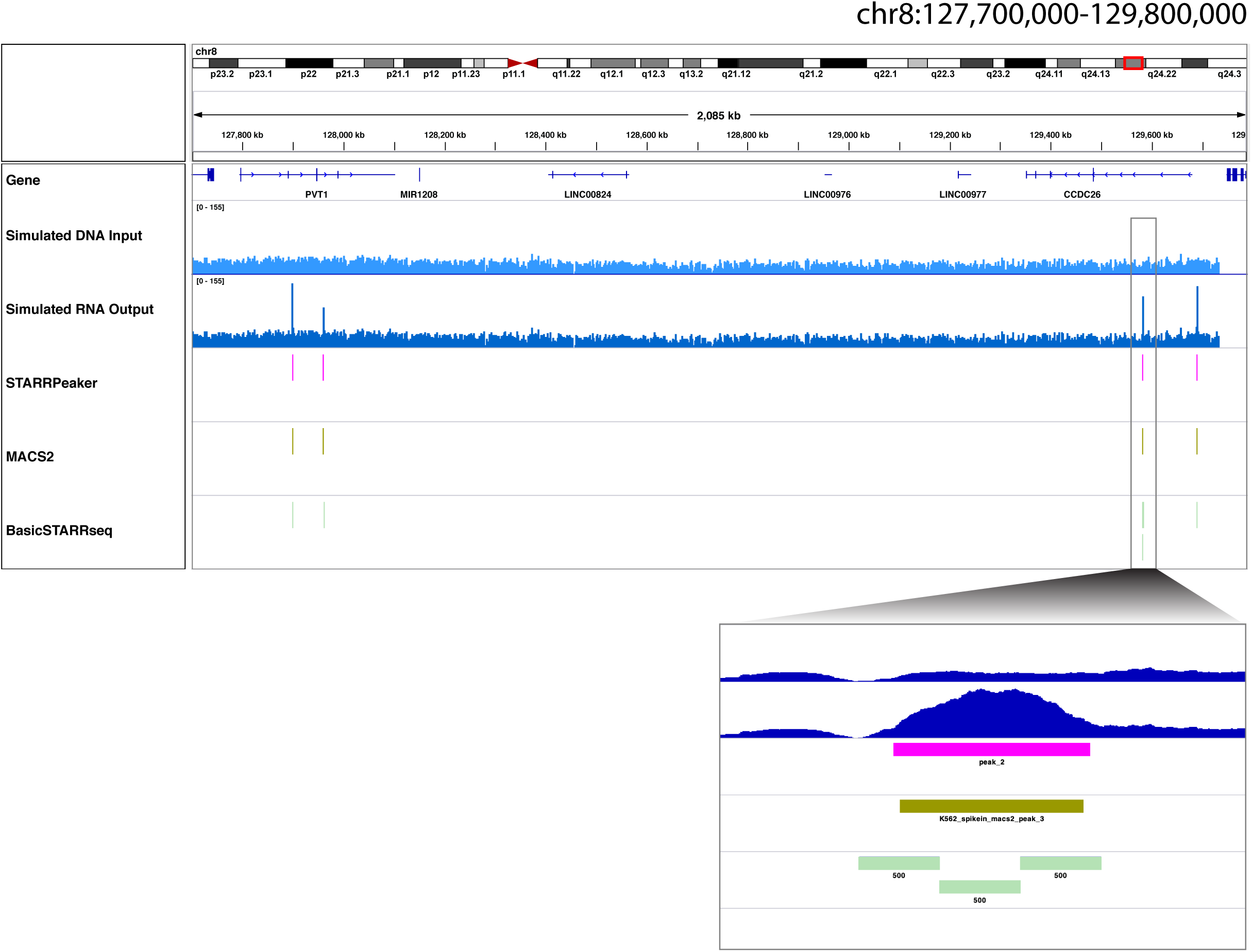
Comparison of peaks identified by various methods using a simulated STARR-seq dataset containing four spike-in control regions.

**Supplementary Figure 7.**
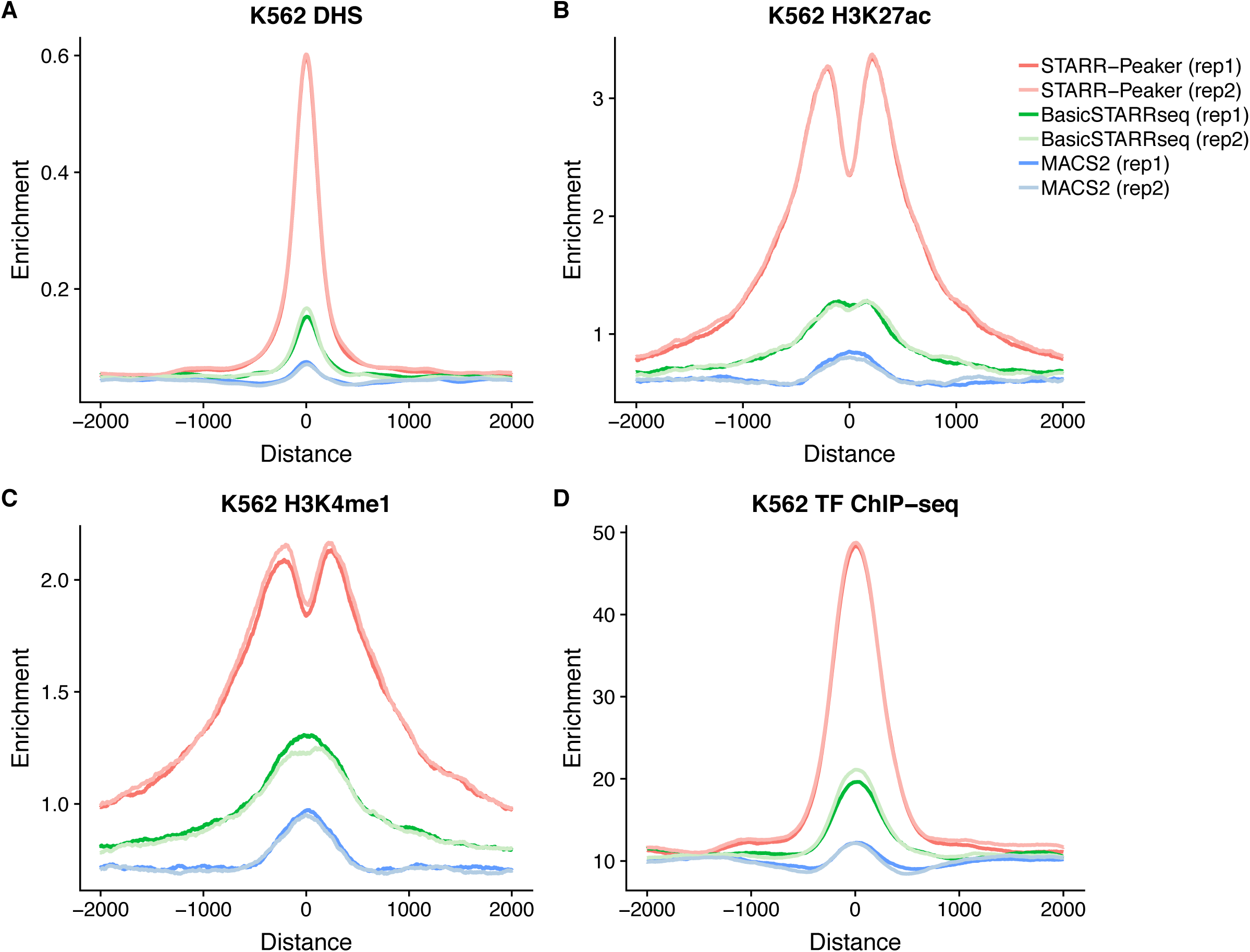
Enrichment of epigenetic signals around peaks in K562. All peaks were centered at the summit, uniformly thresholded using P-value < 0.001, and 10,000 peaks were randomly selected. Aggregated read depth at 2,000 bp upstream and downstream were plotted for **(A)** DNase I hypersensitive sites (DHS), **(B)** H3K27ac, **(C)** H3K4me1, and **(D)** aggregated TF ChIP-seq profile. For DNase-seq, enrichment indicates unique read depth. For histone ChIP-seq, enrichment indicates fold change over control. For TF ChIP-seq aggregate, enrichment indicates the number of TFs binding.

**Supplementary Figure 8.**
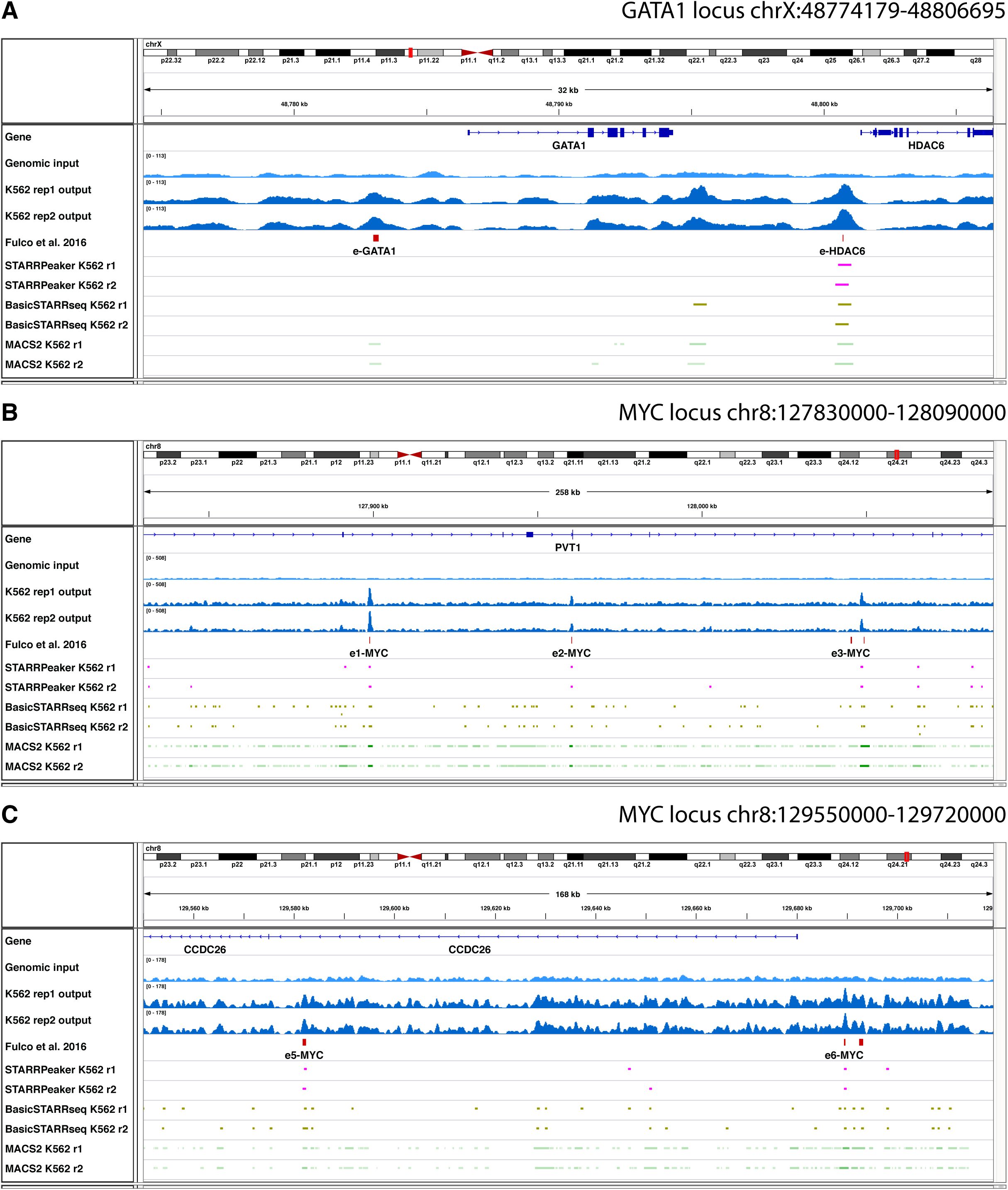
**(A-C)** Genome browser session comparing STARRPeaker to other peak-calling methods at validated enhancers from CRISPRi.

**Supplementary Figure 9.**
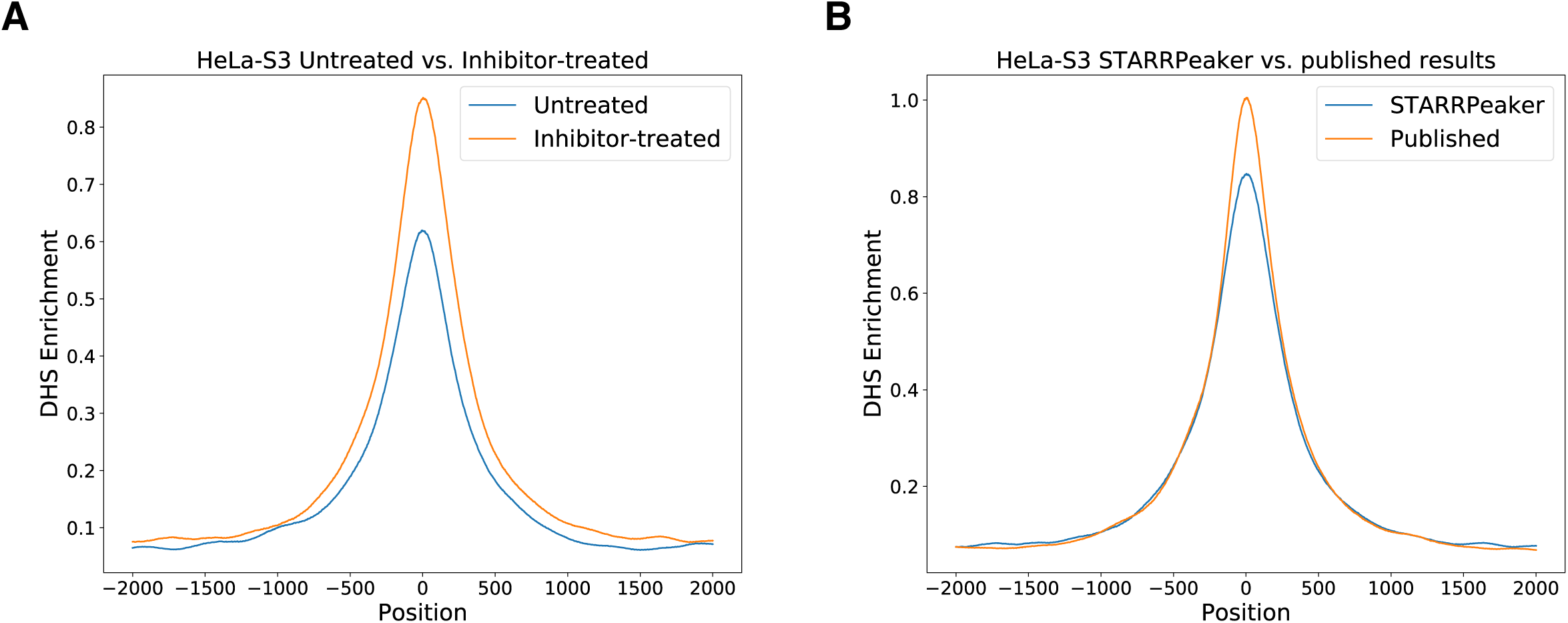
Application of STARRPeaker on an external HeLa-S3 dataset. **(A)** Comparison of chromatin accessibility (DNase-seq) for STARRPeaker peaks between untreated and inhibitor-treated samples. **(B)** Comparison of STARRPeaker peaks to published results. STARRPeaker found 6,540 additional peaks that are enriched with chromatin accessibility signals from a HeLa-S3 sample.

**Supplementary Figure 10.**
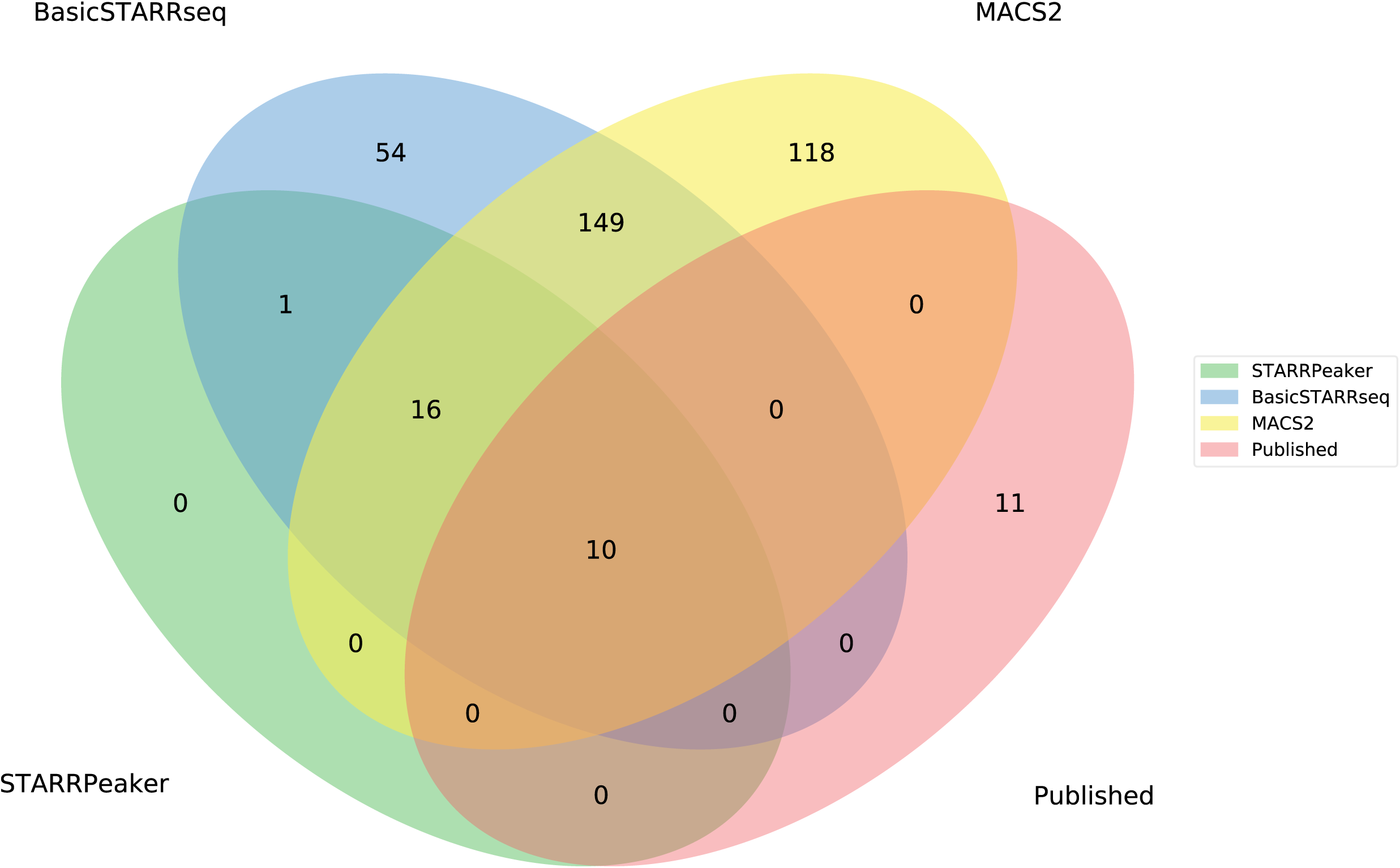
Venn diagram for four-way comparison of peaks identified by various methods using a published dataset from Rathert et al. 2015.

